# A kinetic modeling platform for predicting the efficacy of siRNA formulations *in vitro* and *in vivo*

**DOI:** 10.1101/2022.06.01.494194

**Authors:** Esther H. Roh, Millicent O. Sullivan, Thomas H. Epps

## Abstract

We present a computational modeling protocol that can accurately predict changes in both *in vitro* and *in vivo* gene expression levels in response to the application of various siRNA formulations. Users can obtain crucial information (*i*.*e*., maximum silencing level; duration of silencing) towards the design of therapeutically relevant dosing regimens with experimental measurements from a single time point as an input. This ability to simulate numerous experimental gene silencing scenarios has not been demonstrated previously with other RNA interference models.

**Graphical abstract:** 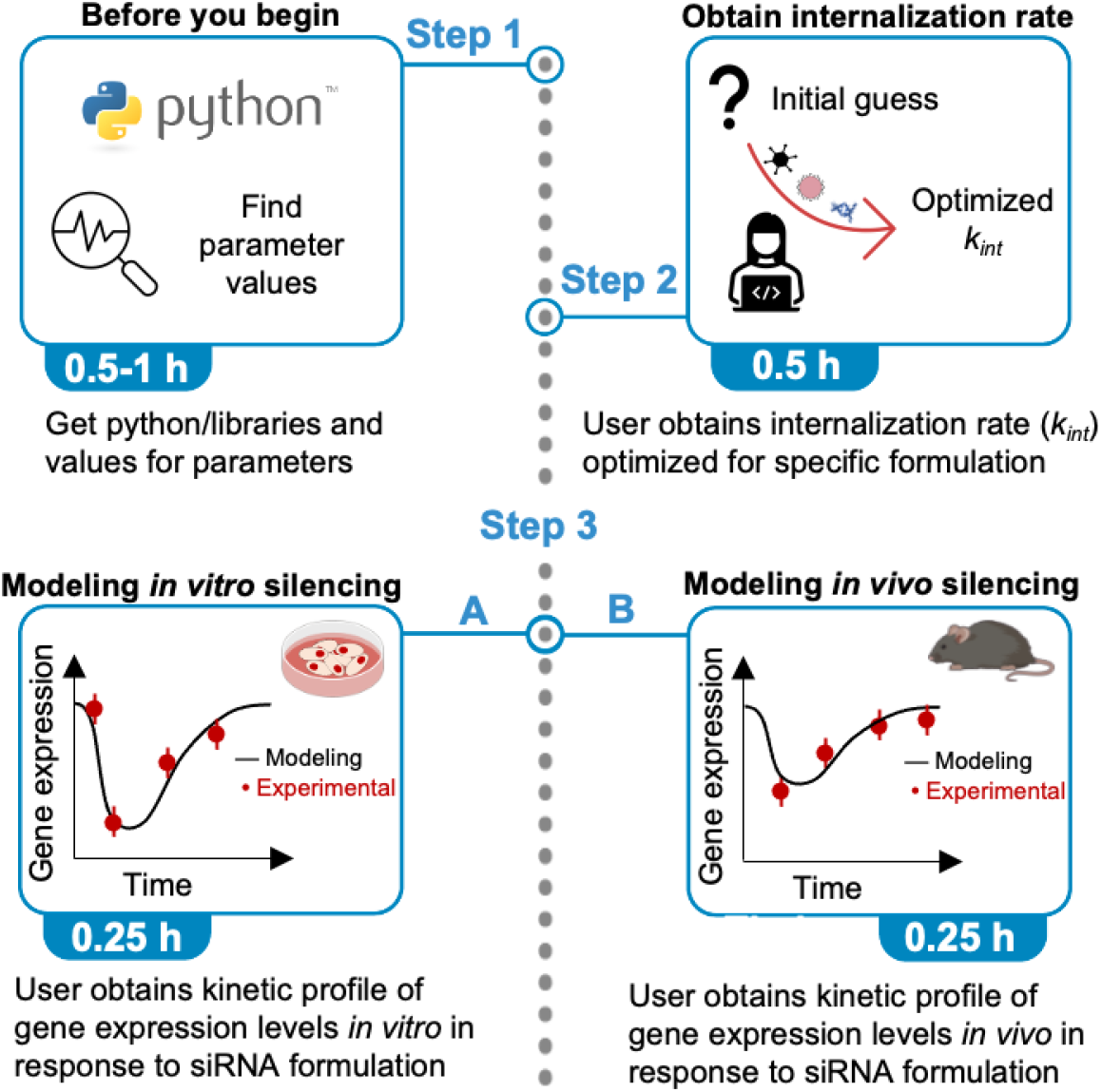

## Before you begin

The protocol below describes the specific steps and values necessary to predict green fluorescent protein (GFP) silencing in B16F10 cells using electrotransfection *in vitro* and apolipoprotein B (apoB) silencing in mouse models using siRNA-lipid nanoparticles. However, we have also used this protocol for various scenarios with different vectors (*e*.*g*., naked siRNA, jetPRIME^®^, Lipofectamine™, Oligofectamine™, chitosan, polymer conjugates, photo-responsive polymers, multifunctional envelope-type nanodevices, and calcium phosphate nanoparticles), different target genes (*e*.*g*., Luciferase, GFP, glyceraldehyde 3-phosphate dehydrogenase, interleukin 1 beta, voltage dependent anion channel 1, PTEN), and different cell types (*e*.*g*., B16F10, Neuro2A, HeLa, LNCaP, H1299, A549).

### Downloading and the launching Python platform

#### Timing: 0.5 – 1 h

This section includes the minimal hardware requirements, software requirements, and input parameters needed to run the model.

#### Hardware

Personal computer with a minimum of 5 GB free disk space and 32 GB random-access memory.

### Software requirements

A convenient method to run Python on any operating system is to use Anaconda, an open-source distribution for Python and R for scientific computing.

1. Anaconda can be downloaded from https://www.anaconda.com/ according to individual computer specifications.
2. Once Anaconda is installed, open the Anaconda Navigator, click on the ‘Environments’ tab to the left, and make sure the following Python packages are installed: numpy (Harris et al., 2020), scipy (Virtanen et al., 2020), sklearn.metrics (Pedregosa et al., 2011), and matplotlib (Hunter, 2007). All of the packages mentioned should be pre-installed in the base environment.
3. Click on the ‘Home’ tab to the left and launch Jupyter Notebook.

**Figure 1.**
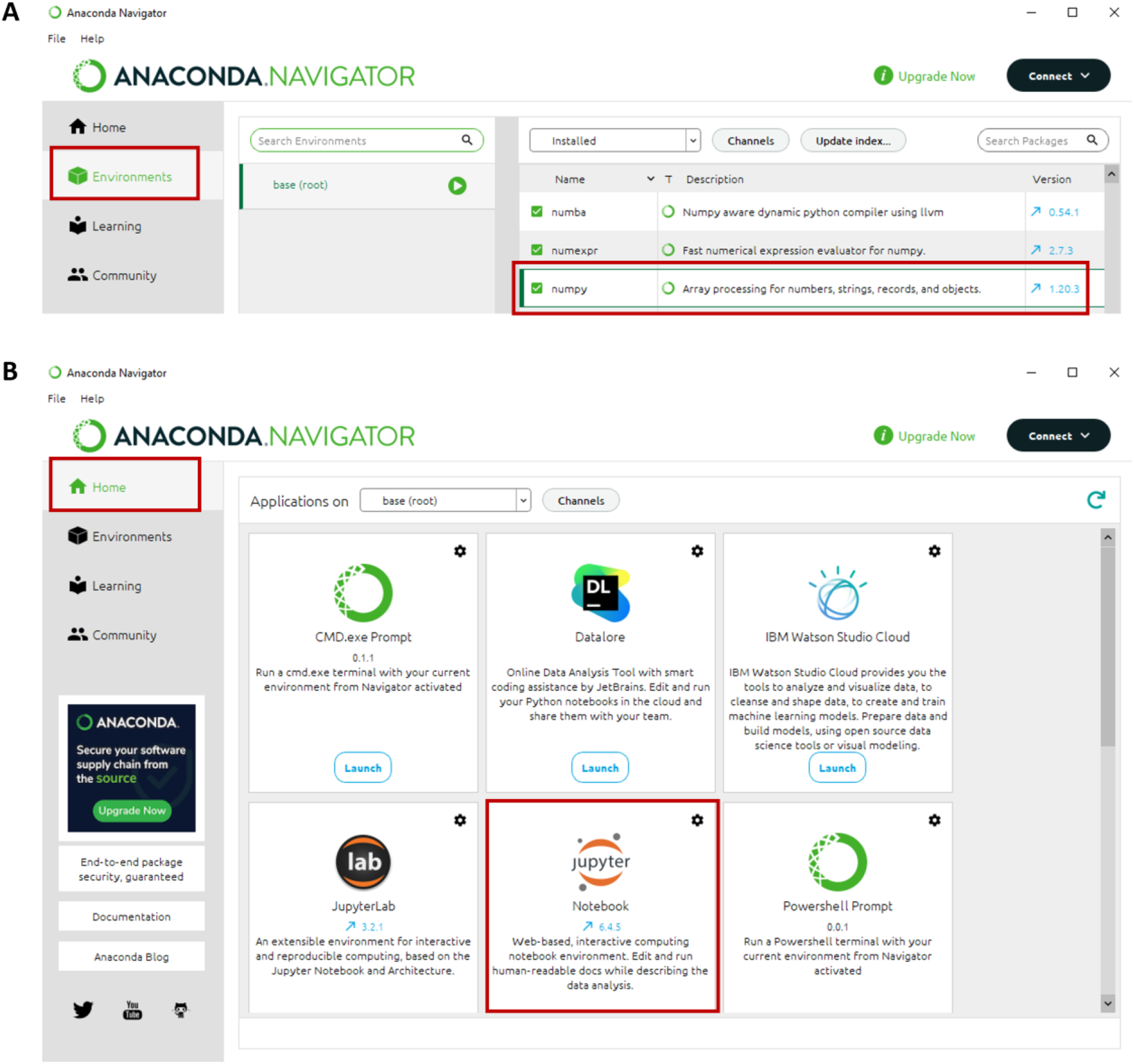
Launching Anaconda and preparing required Python packages. (A) After launching Anaconda, make sure that the appropriate Python packages are available in the ‘Environments’ tab. (B) Once all packages are available, launch Jupyter Notebook in the ‘Home’ tab.

### Preparing input parameters

#### Timing: 0.5 – 72 h

To employ the kinetic model provided in this protocol, users are required to provide the values for the following parameters.

A. siRNA half-life
B. mRNA half-life
C. protein half-life
D. cell doubling time
E. transfection volume
F. transfection time
G. transfection concentration
H. *in vitro* experimental measurements of gene expression levels at one or more time points, preferably between 48 – 72 h

***Note:*** There are numerous protocols available from either commercial websites (*e*.*g*., ThermoFisher, Dharmacon) or peer-reviewed literature (Sakurai et al., 2010) that detail steps for obtaining this type of experimental measurement

## Key resources table

| REAGENT or RESOURCE | SOURCE | IDENTIFIER |
| --- | --- | --- |
| Software and algorithms |  |  |
| Fitting internalization rate | This paper | N/A |
| *In vitro* model | This paper | N/A |
| *In vivo* model | This paper | N/A |
| Other |  |  |
| Internalization rate of external siRNA ($k_{int}$) | This paper | N/A |
| Transfection volume ($V_{ex}$) | Specified experimentally | N/A |
| Intracellular volume ($V_{in}$) | (Bartlett and Davis, 2006, Wittrup et al., 2015) | $V_i$ |
| Dilution of siRNA due to cell division ($k_{s,dil}$) | Calculated from cell doubling time | N/A |
| Intracellular degradation of siRNA ($k_{s,deg}$) | (Bartlett and Davis, 2006) | $k_{deginna}$ |
| Intracellular RISC concentration ($RISC_{tot}$) | (Bartlett and Davis, 2006) | $rtot$ |
| Formation of active RISC ($k_{RISC}$) | (Bartlett and Davis, 2006) | $k_{formRISC}$ |
| Intracellular RISC degradation ($k_{r,deg}$) | (Bartlett and Davis, 2006) | $k_{degRISC}$ |
| Intracellular mRNA production rate ($k_{mRNA}$) | Calculated from mRNA half-life | N/A |
| Intracellular mRNA degradation rate ($k_{m,deg}$) | Calculated from mRNA half-life | N/A |
| Cleavage rate of mRNA ($k_{cleav}$) | (Roh et al., 2021) | $k_{cleav}$ |
| Intracellular protein production rate ($k_{prot}$) | Calculated from protein half-life | N/A |
| Intracellular protein degradation rate ($k_{p,deg}$) | Calculated from protein half-life | N/A |

## Step-by-step method details

### Fitting for internalization rate

#### Timing: 15 – 30 min

**CRITICAL**: To proceed with the following steps, make sure to first install all software and packages listed in ‘Software requirements.’

One of the core aims of our protocol is to provide users with the ability to easily predict changes in gene expression levels both *in vitro* and *in vivo* upon application of any siRNA formulation of interest. Traditionally, modeling the efficacy of various siRNA formulations is challenging because the RNAi process consists of multiple steps that are difficult to characterize experimentally.(Haley and Zamore, 2004, Cuccato et al., 2011, Bartlett and Davis, 2006) In particular, cellular uptake and necessary post-uptake processing, including endosomal escape, comprises multiple steps, and the parameters that describe these steps are unique to each delivery system. In this mathematical framework, we define the internalization rate as a single aggregated parameter so that this rate is the only unknown variable, and thus, can be fit from a single experimental time point. By fitting for an aggregated internalization rate, users can reduce the number of parameters needed to implement the model in various experimental scenarios. This fitting process is performed as follows:

1. To call the packages that will be used for this protocol, copy and paste the following code into Jupyter Notebook.

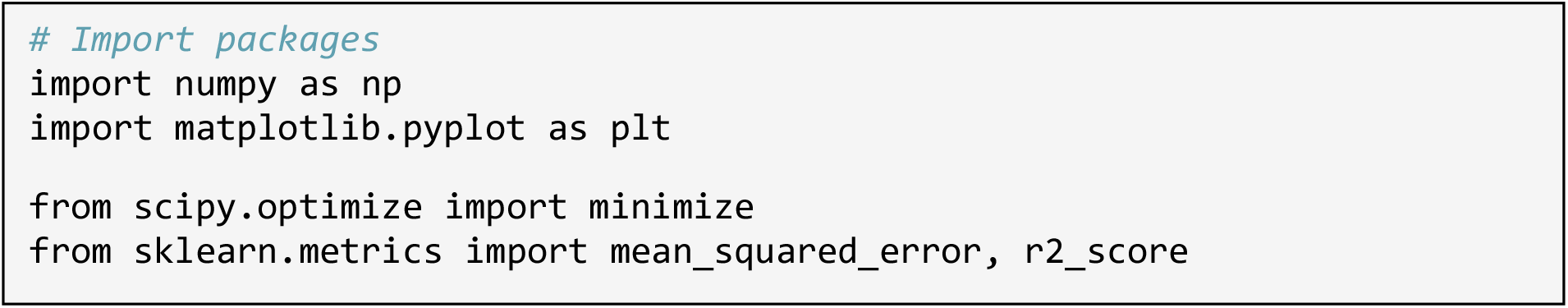
2. Copy and paste the following code and modify the values for the seven variables accordingly.

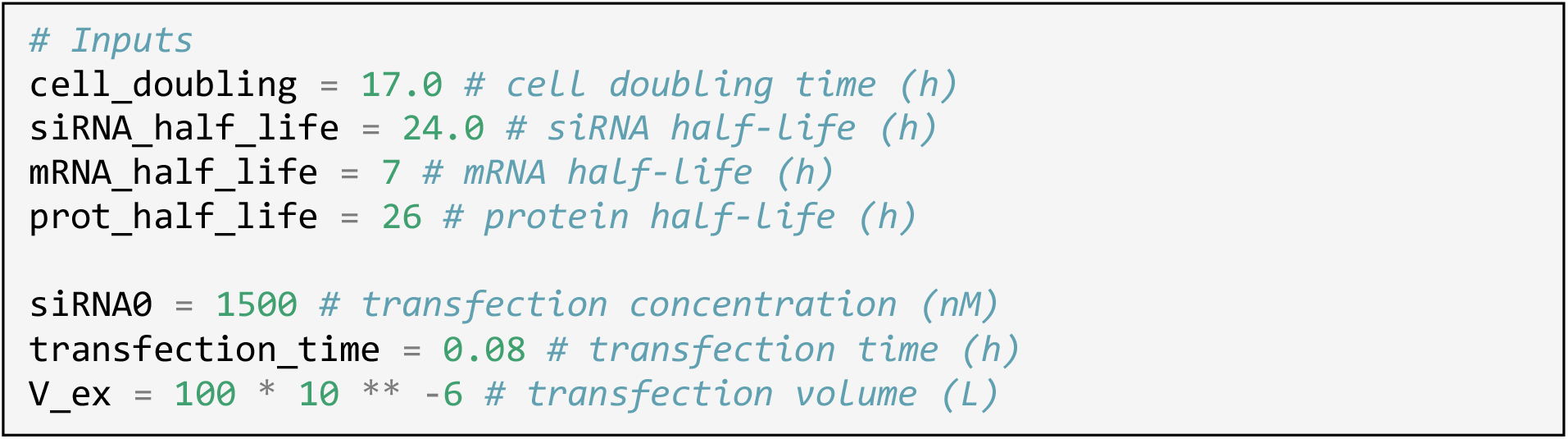 ***Note:*** Cell-doubling rates are needed to correctly account for the dilution of intracellular siRNA over time. The next three parameters are required to compute the component degradation rates and the production rates of mRNA and protein assuming steady-state values in the absence of siRNA. The last three parameters are needed to correctly scale and account for the initial siRNA concentration available to the cells.
3. Copy and paste the following code, and define the final time point of interest as the variable “tf_transfection”.

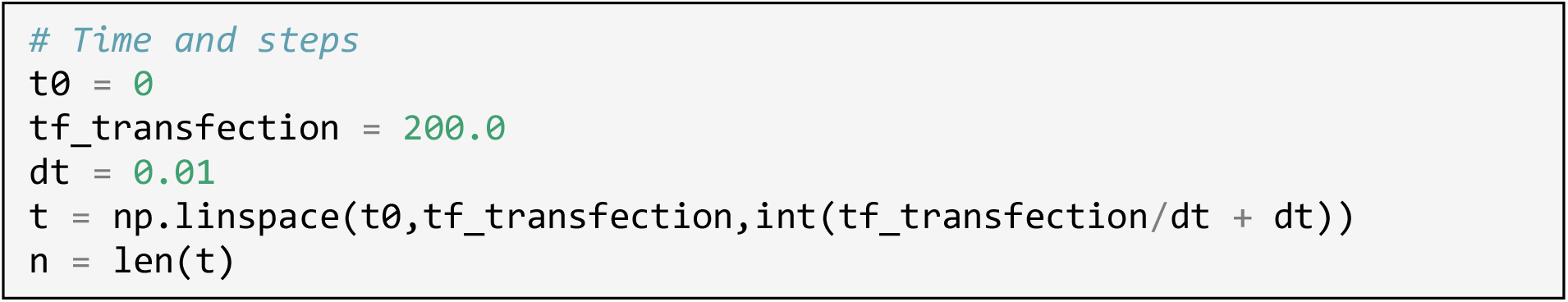 ***Note:*** The accuracy of model predictions is enhanced as more steps are taken. However, increasing the number of steps results in an escalation of computation cost (*e*.*g*., time). A minimum value of 0.01 is recommended based on characteristic time calculations of the fastest rate (cleavage rate of mRNA = 500 nM^-1^h^-1^). (Roh et al., 2021, Wittrup et al., 2015)
4. Copy and paste the following code, and input the time points of experimental data as parameters of variable “x”, and input the respective gene expression levels at each time point as parameters of variable “y”.

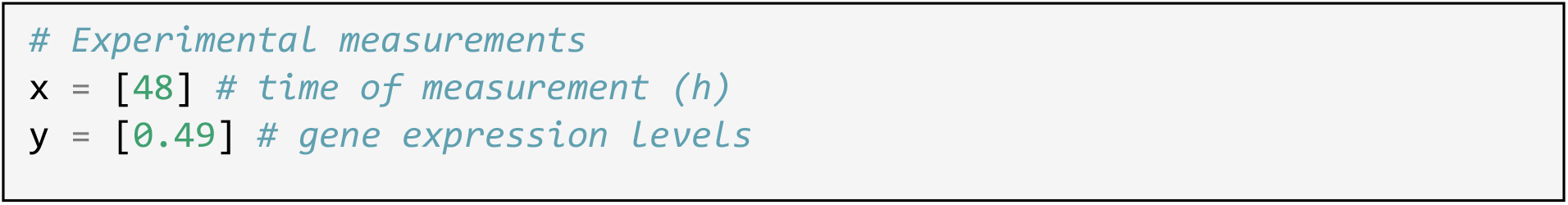 ***Note:*** Data for y-array are defined as protein expression levels post-siRNA exposure that is scaled between the values of 0 and 1.
5. Copy and paste the following code, and input an initial guess for internalization rate.

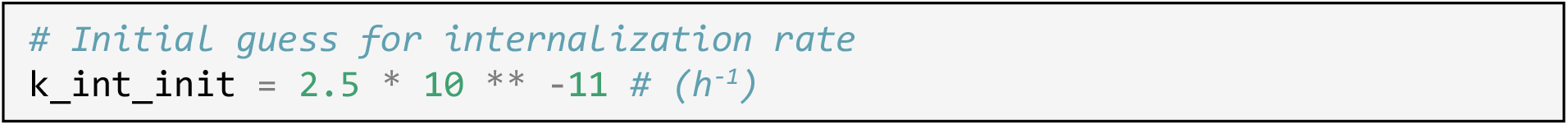 ***Note:*** The recommended starting value is 10^−10^ h^-1^.
6. Copy and paste the following code, run code, and obtain an optimized internalization rate.

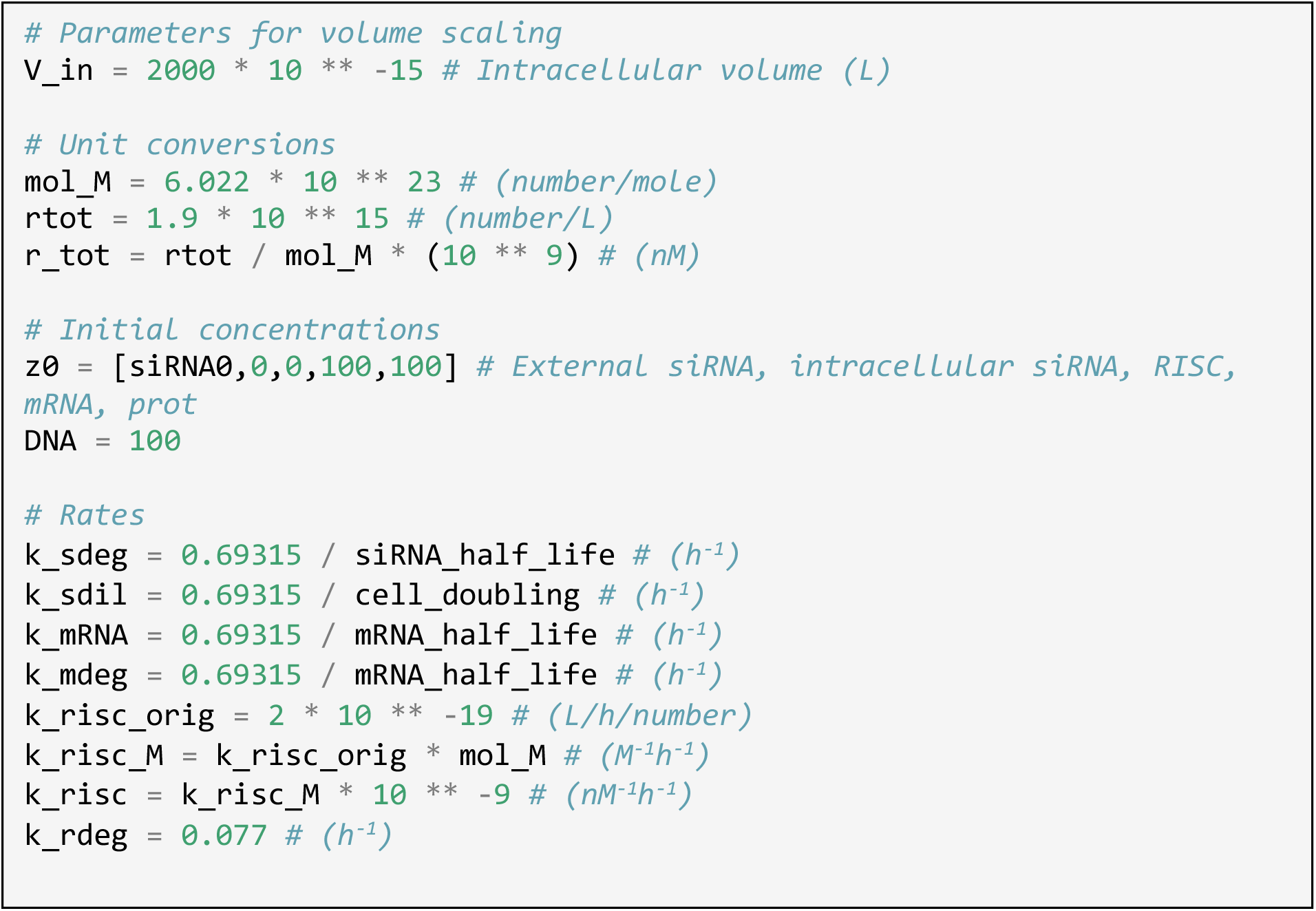
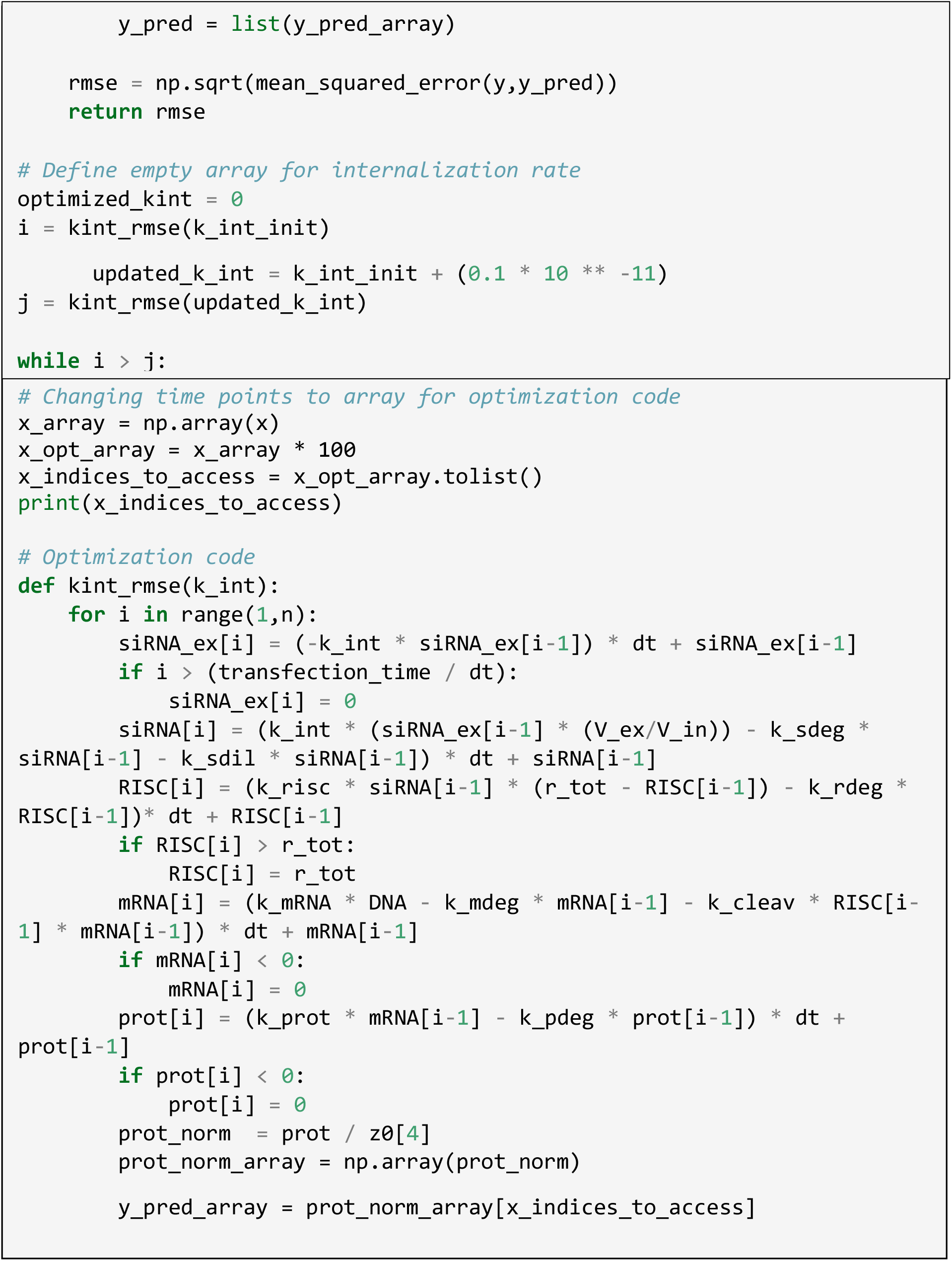 ***Note:*** Do not change any values within the code provided in step 6. At the end of the run, the code will automatically report the optimized internalization rate along with the root-mean-square error (RMSE) associated with that rate.

**Figure 2.**
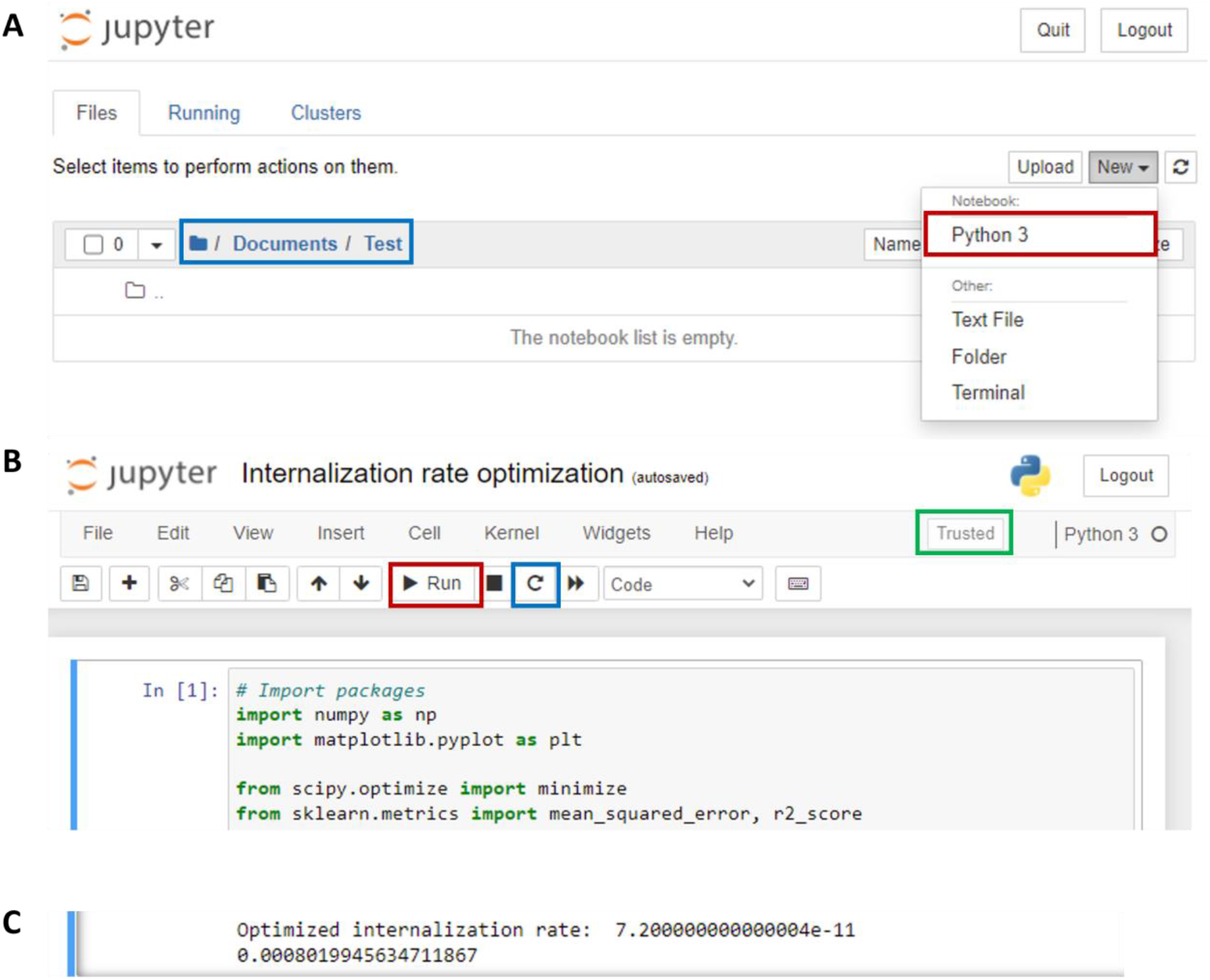
Fitting for an optimized internalization rate. (A) After opening Jupyter Notebook, choose the correct directory in which to save your notebook (blue box). Click on the ‘New’ box and create a new Python 3 notebook (red box). (B) After copy and pasting all of the code provided in the ‘Fitting for internalization rate’ section, run code (red box). If there is an error and users must re-run the code for any reason, make sure to reset the kernel (blue box) before re-running. Make sure that the kernel is trusted (green box). (C) Once the code is finished, the optimized internalization rate and RMSE for that rate is displayed.

### Modeling siRNA-induced changes in gene expression levels *in vitro*

#### Timing: 15 min

**CRITICAL**: To proceed with the following steps, make sure to first obtain an optimized internalization rate using the procedure listed in ‘Fitting for internalization rate.’

To expand the applicability of the model described in this protocol, the number of parameters required to accurately predict gene silencing *in vitro* following the delivery of an siRNA formulation is further reduced by including only the most important parameters within the mathematical framework to get quantitative agreement with experimental data. Major rate-limiting kinetic steps for the four main components of the RNAi process (*i*.*e*., siRNA, RNA-induced silencing complex (RISC), mRNA, and protein) are identified from literature and incorporated into the model. In the following paragraph, the parameters that are considered for each component, along with the rationale for choosing those specific kinetic rates for *in vitro* modeling of siRNA-induced gene silencing, are detailed.

**Table 2.** Model variables.

| **Variable (Name)** | **Description (Units)** |
| --- | --- |
| **siRNA<sub>ex</sub>** | External siRNA (nM) |
| **siRNA<sub>in</sub>** | Intracellular siRNA (nM) |
| **RISC** | Activated RISC (RISC + siRNA) (nM) |
| **mRNA** | Intracellular mRNA (nM) |
| **DNA** | Intracellular DNA (nM) |
| **prot** | Intracellular protein (nM) |

**Table 3.** Model parameters and values.

| Parameter (Name) | Description | Determination | Value (Units) |
| --- | --- | --- | --- |
| $k_{int}$ | Internalization rate of external siRNA | Fit to each system | (h<sup>-1</sup>) |
| $V_{ex}$ | Transfection volume | Specified experimentally | (L) |
| $V_{in}$ | Intracellular volume | (Wittrup et al., 2015, Bartlett and Davis, 2006) | $2 * 10^{-12}$ (L) |
| $k_{s,dil}$ | Dilution of siRNA due to cell division | Calculated from cell doubling time | (h<sup>-1</sup>) |
| $k_{s,deg}$ | Intracellular degradation of siRNA | Calculated from siRNA half-life | (h<sup>-1</sup>) |
| $RISC_{tot}$ | Intracellular RISC concentration | (Bartlett and Davis, 2006) | 3.2 (nM) |
| $k_{RISC}$ | Formation of active RISC | (Bartlett and Davis, 2006) | $1.2 * 10^{-4}$ (nM<sup>-1</sup>h<sup>-1</sup>) |
| $k_{r,deg}$ | Intracellular RISC degradation | (Bartlett and Davis, 2006) | $7.7 * 10^{-2}$ (h<sup>-1</sup>) |
| $k_{mRNA}$ | Intracellular mRNA production rate | Steady-state approximation, calculated from mRNA half-life | (h<sup>-1</sup>) |
| $k_{m,deg}$ | Intracellular mRNA degradation rate | Calculated from mRNA half-life | (h<sup>-1</sup>) |
| $k_{cleav}$ | Cleavage rate of mRNA | (Roh et al., 2021) | $5 * 10^2$ (nM<sup>-1</sup>h<sup>-1</sup>) |
| $k_{prot}$ | Intracellular protein production rate | Steady-state approximation, calculated from protein half-life | (h<sup>-1</sup>) |
| $k_{p,deg}$ | Intracellular protein degradation rate | Calculated from protein half-life | (h<sup>-1</sup>) |

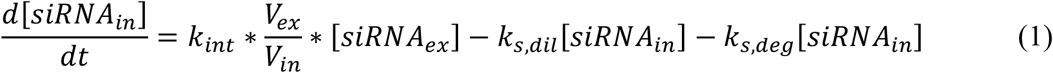

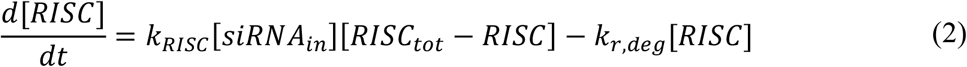

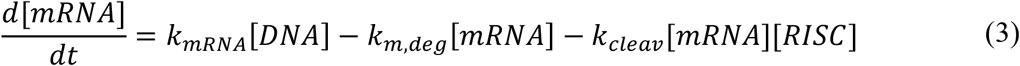

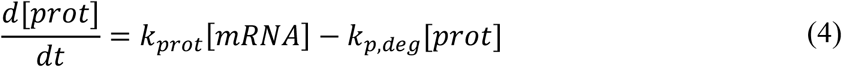

As noted in the previous section, the increase in intracellular siRNA concentration is defined to depend solely on a single aggregated internalization rate (*k*_*int*_, Eq.1). Additionally, the amount of intracellular siRNA is affected by ***(i)*** dilution of the local intracellular siRNA concentration as cells divide (*k*_*s,dil*_, Eq.1); and ***(ii)*** siRNA degradation (*k*_*s,deg*_, Eq.1). To model the RISC concentration, we assume that the concentration of active RISC, *e*.*g*., the species that can recognize and cleave mRNA, depends solely on the binding of free siRNA to RISC (*k*_*RISC*_, Eq.2) and RISC degradation (*k*_*r,deg*_, Eq.2). Although previous studies have shown that RISC has multiple intermediates,(Pratt and MacRae, 2009) our assumption is reasonable because the formation of the siRNA-RISC intermediate has the slowest kinetic rate of all steps in active RISC formation.(Pratt and MacRae, 2009) Meanwhile, the mRNA concentration is modeled to depend on innate mRNA production (*k*_*mRNA*_, Eq.3), degradation (*k*_*m,deg*_, Eq.3), and cleavage (*k*_*cleav*_, Eq.3) rates. Finally, protein concentration depends directly on protein production from the mRNA template (*k*_*prot*_, Eq.4), along with protein degradation (*k*_*p,deg*_, Eq.4).

7. To call the packages that will be used for this protocol, copy and paste the code provided below into Jupyter Notebook.

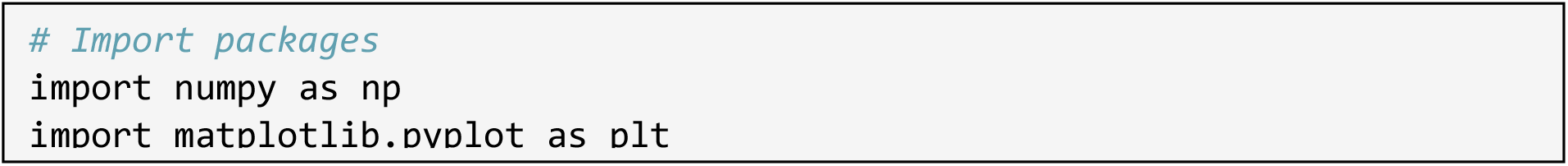
8. Copy and paste the following code, and modify the values for the seven variables accordingly.

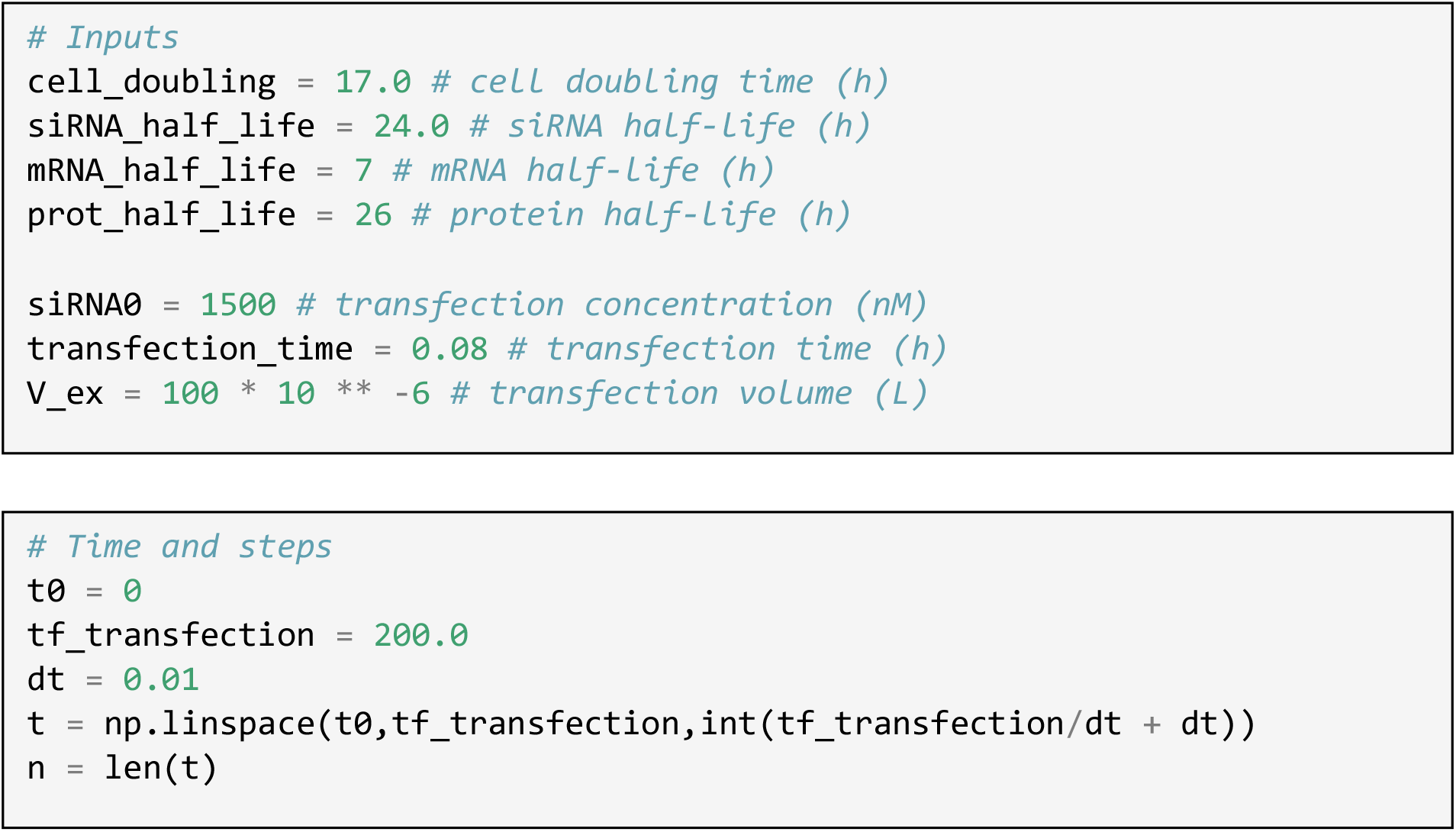
9. Copy and paste the following code, and define the final time point of interest as the variable “tf_transfection”.
10. Copy and paste the following code, and input the internalization rate fitted using the previous procedure.
11. Copy and paste the following code, run code, and obtain changes in mRNA and protein concentration levels.

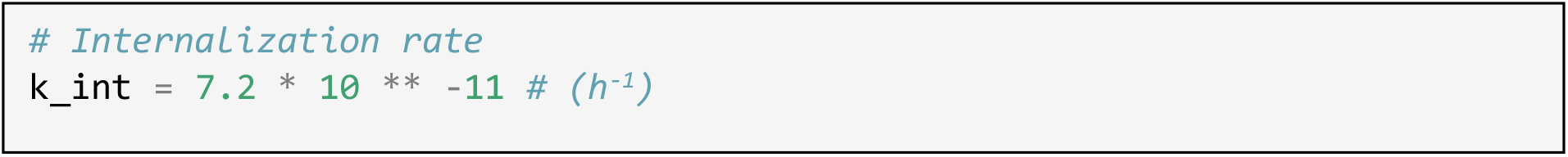
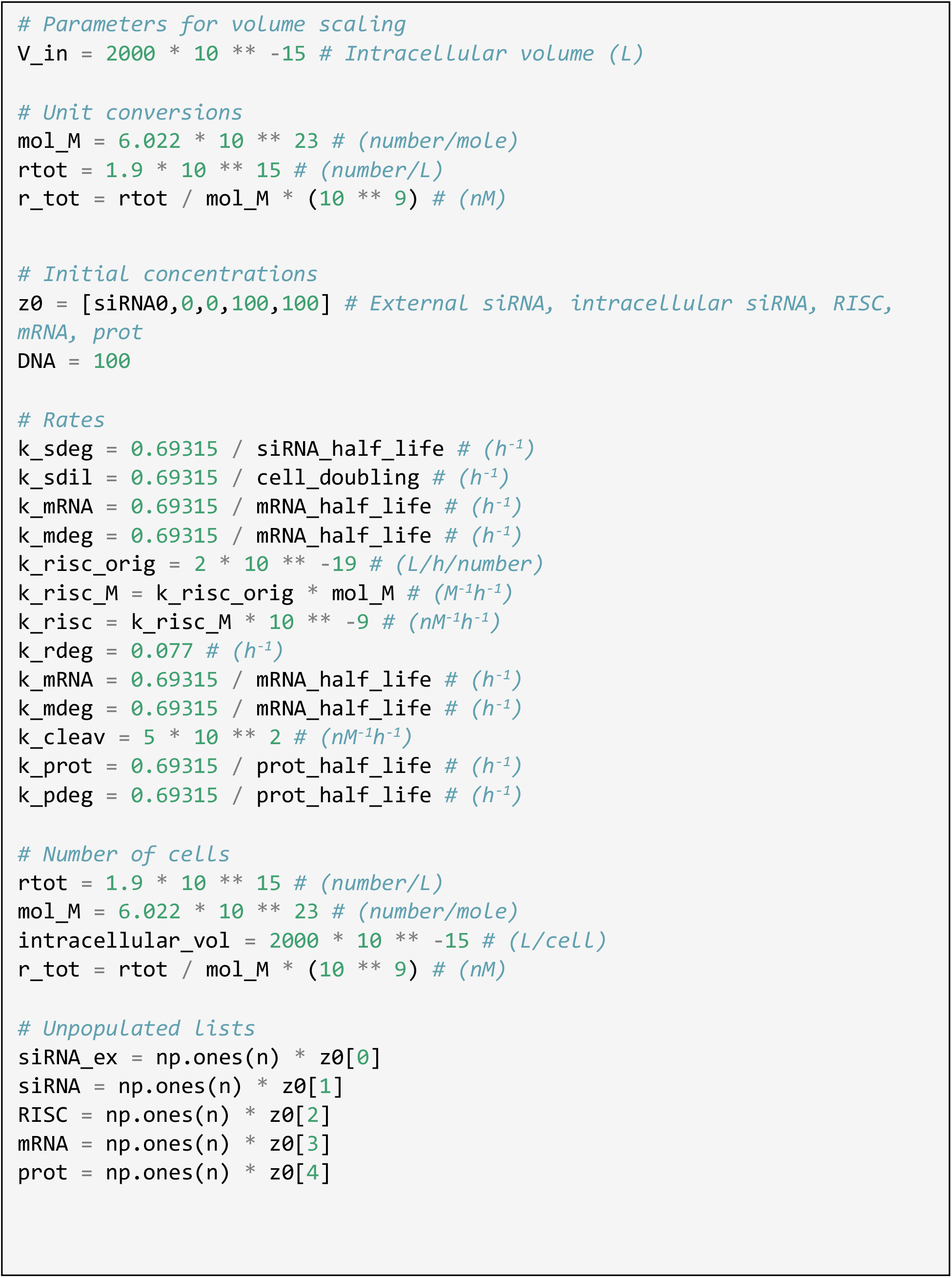
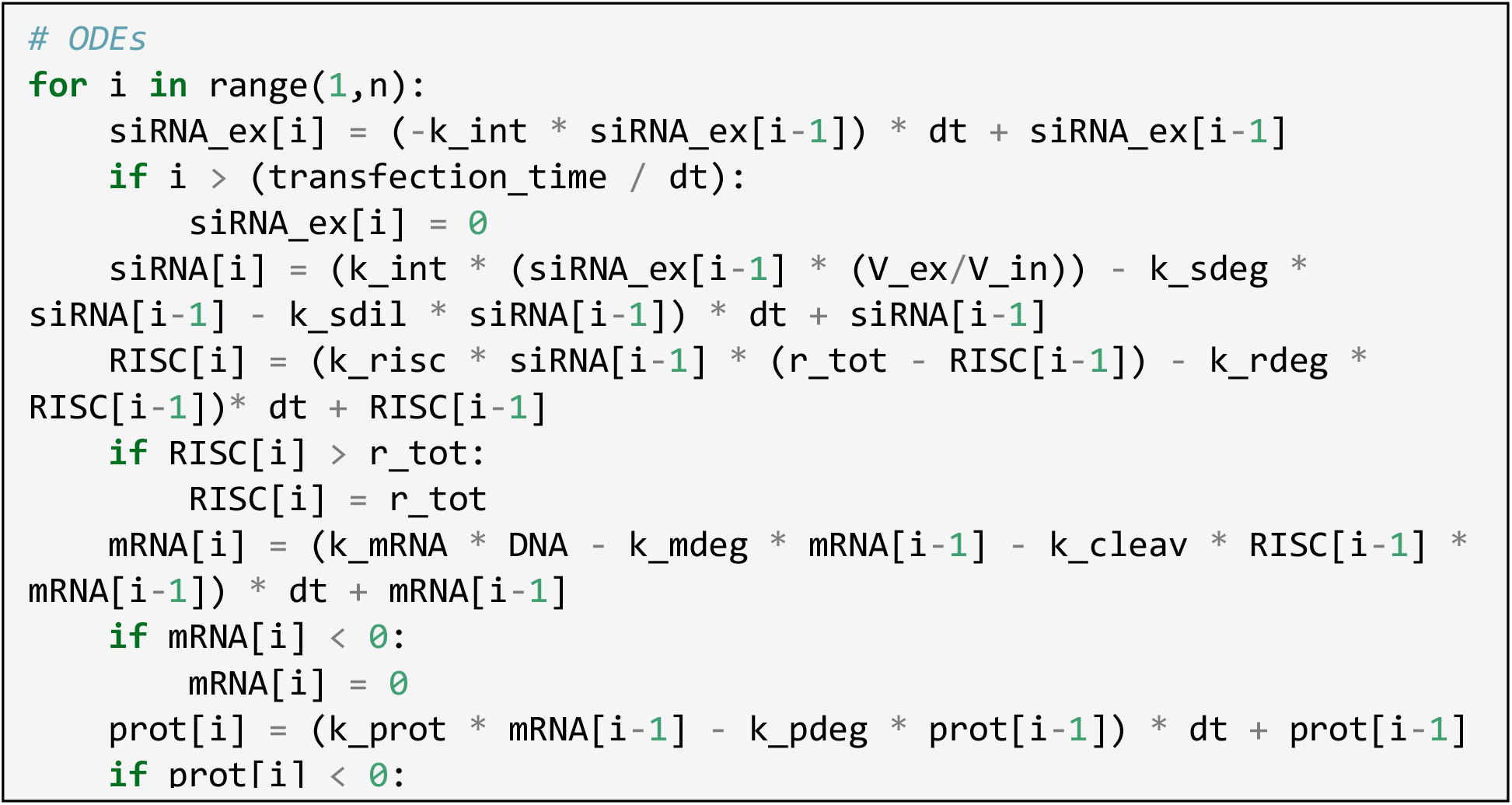
12. Copy and paste the following code, and plot desired data accordingly.

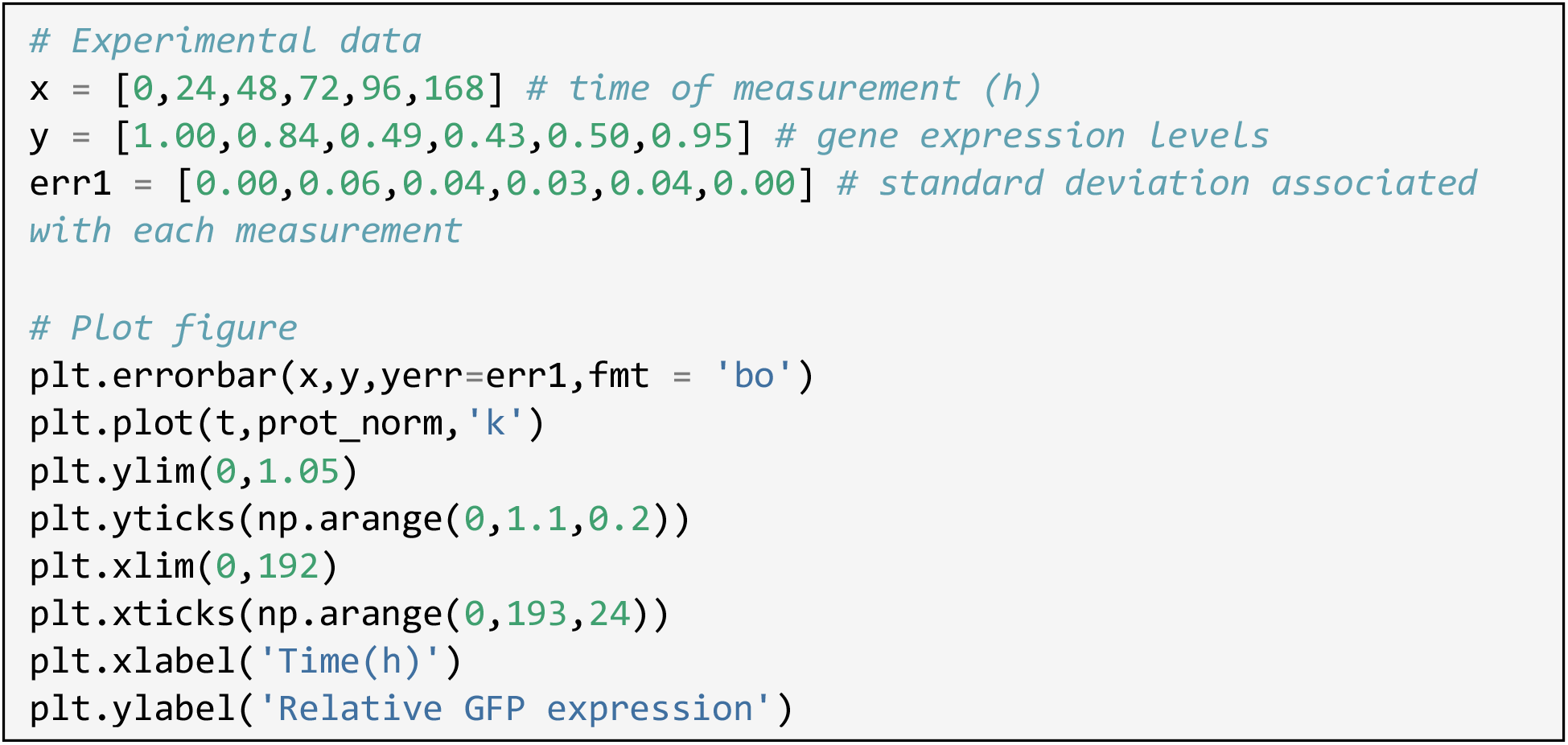 ***Note:*** In this example code, experimental data regarding GFP expression levels over time are incorporated into the code. Users can change data within the x-, y- and err1-arrays for time points, gene expression levels, and associated error, respectively. The ‘# Experimental data’ section of the code can be omitted if users do not wish to plot experimental and modeling data together. Also, plot details including plot type, x-axis and y-axis ticks, x-axis and y-axis limits can be customized to meet the users’ needs. Detailed explanations and examples can be found at matplotlib.org.(Hunter, 2007)

**Figure 3.**
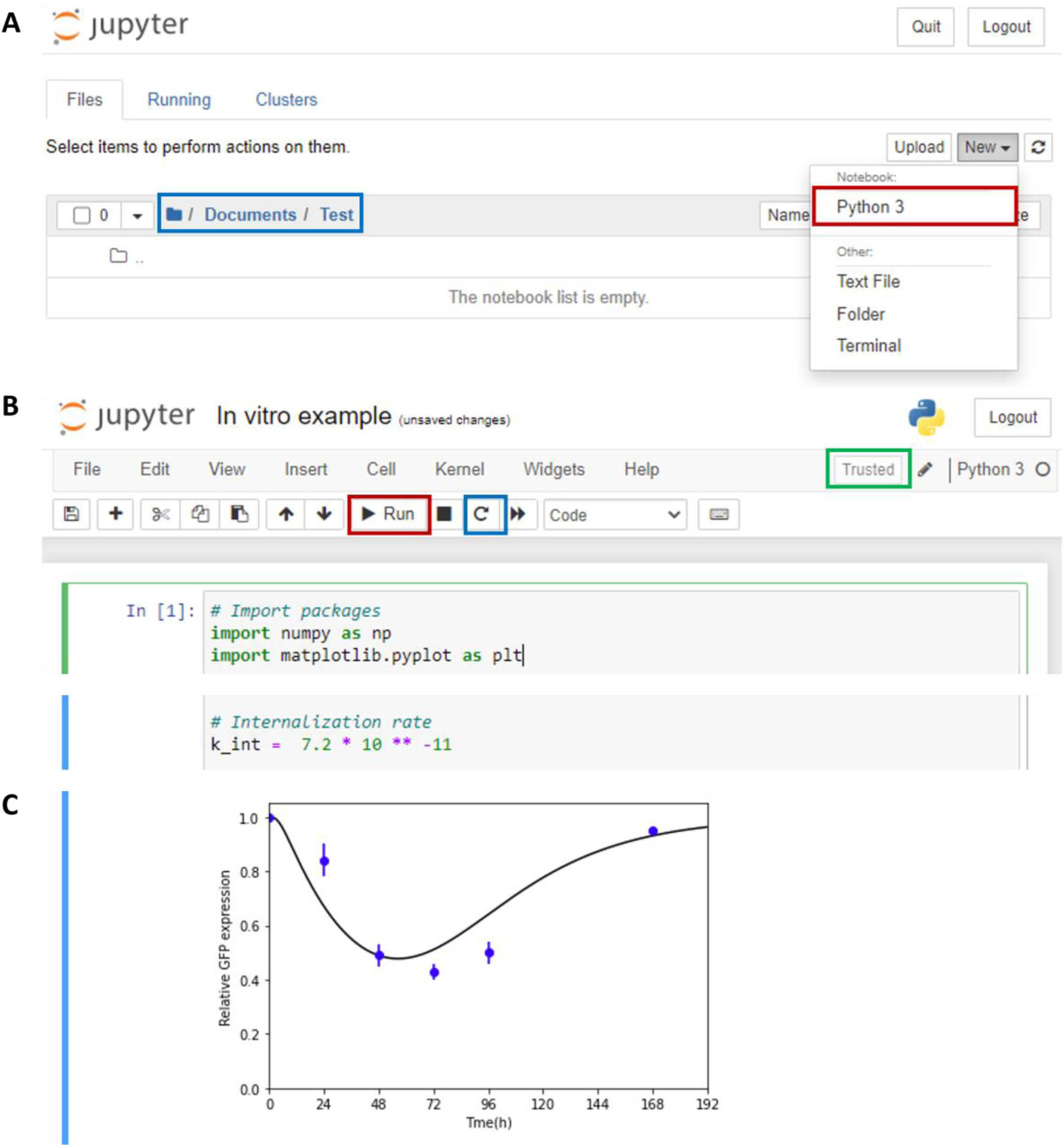
Modeling changes in gene expression levels *in vitro* using the fitted internalization rate. (A) As mentioned in previous section, after opening Jupyter Notebook, choose the correct directory in which to save your notebook (blue box). Click on the ‘New’ box and create a new Python 3 notebook (red box). (B) Copy and paste all code provided in the ‘Modeling siRNA-induced changes in gene expression levels *in vitro*’ section. Make sure that the internalization rate is the same as the optimized rate obtained in the ‘Fitting for internalization rate’ section. If there is an error and users must re-run the code for any reason, make sure to reset the kernel (blue box) before re-running. Make sure that the kernel is trusted (green box). (C) Once the code is finished, a plot will be generated with the details and data provided in the ‘# Plot figure’ section of the code.

### Modeling siRNA-induced changes in gene expression levels *in vivo*

#### Timing: 15 min

**CRITICAL**: To proceed with the following steps, make sure to first obtain an optimized internalization rate using the procedure listed in ‘Fitting for internalization rate.’

A major difference between biological parameters that directly affect intracellular siRNA concentrations *in vitro vs. in vivo* is that the clearance of nucleic acids at the target tissue *in vivo* results in siRNA dilution rates that are significantly different from the dilution rates *in vitro*.(Oraiopoulou et al., 2017) More specifically, the dominating siRNA dilution factor *in vitro* is cell division. However, siRNA dilution *in vivo* is governed by additional extracellular delivery barriers. Additional terms were added to the mathematical framework to account for this difference in extracellular siRNA concentration prior to cellular processing. Again, parameters were lumped into two process-related terms – accumulation and clearance from the tissue of interest – to eliminate the need to characterize a series of hard-to-measure transport steps.(Blanco et al., 2015) The accumulation of siRNA in the tissue of interest was defined as a constant and approximated from fractional biodistribution values characterized in the literature,(Hoshyar et al., 2016, Jasinski et al., 2018, He et al., 2010, Pérez-Campaña et al., 2013) because transport steps before accumulation into the target tissue, such as clearance from blood, occur rapidly and can be assumed to be instantaneous in comparison to the rates of later steps.(Alexis et al., 2008) Particle clearance rates from specific tissues are not as widely characterized as accumulation percentages, and users are required to vary the clearance rate to obtain an accurate kinetic profile for *in vivo* gene silencing.

13. To call the packages that will be used for this protocol, copy and paste the code provided below into Jupyter Notebook.

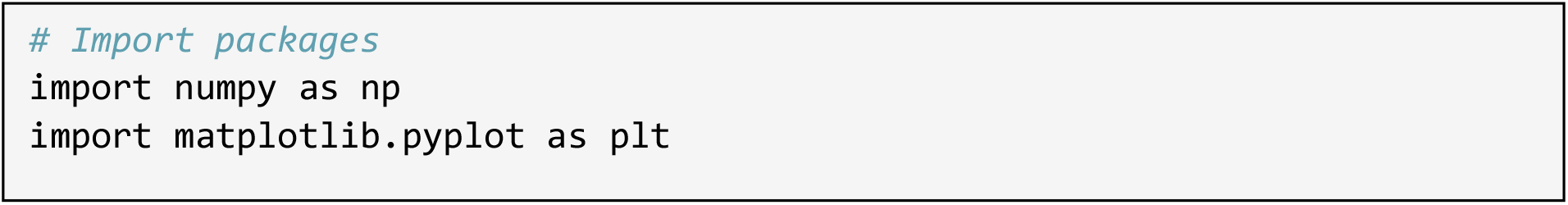
14. Copy and paste the code below, and modify the values for the eight variables accordingly. Estimate the percentage of particles that arrive at the tissue of interest (defined as the variable

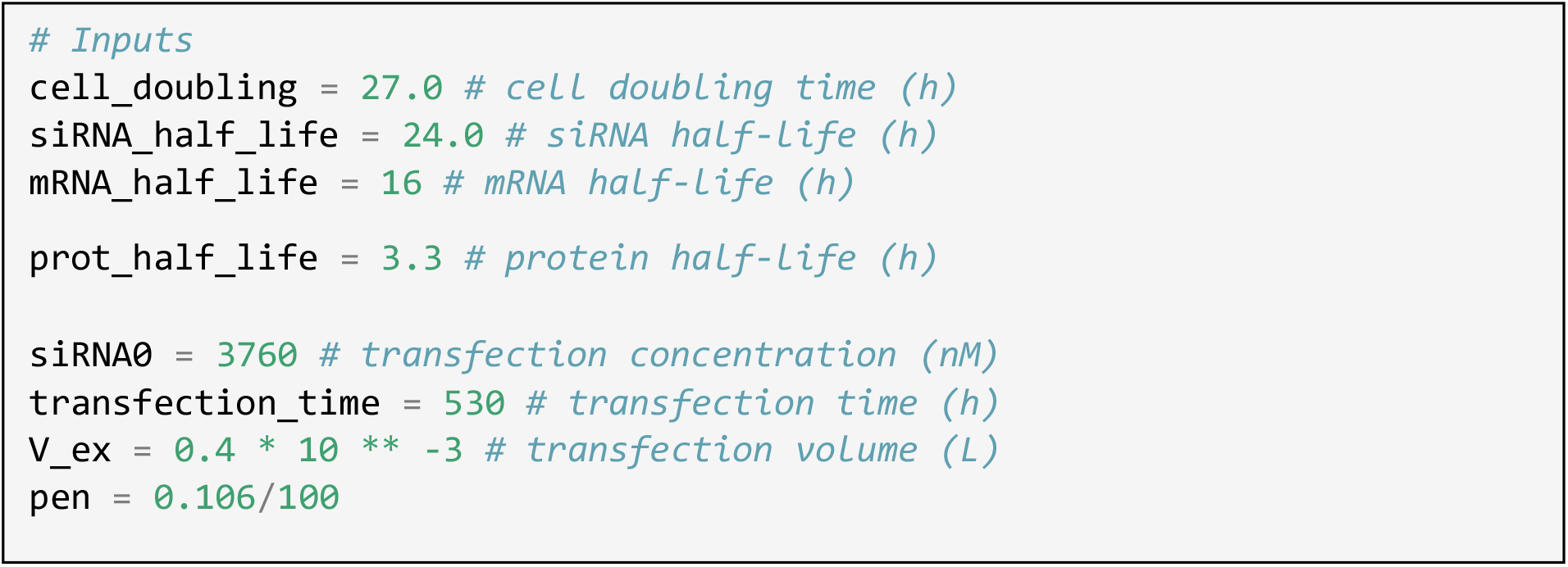 “pen”) using the size and material of nanocarriers from existing literature. ***Note:*** Users can directly input the seven parameters used in the ‘Modeling siRNA-induced changes in gene expression levels *in vitro*’ section IF the *in vitro* data set uses the same siRNA formulation to target the same cell type. The example code given here includes different parameter values from the previous section because the *in vivo* example is modeling a different experimental scenario. ***Note:*** Accumulation (“pen”) is defined as the fractional biodistribution within the tissue of interest, ranging between values of 0 and 1, and is needed to accurately account for the initial external siRNA concentration that is available for cellular internalization. Users are encouraged to estimate this value from literature on the basis of particle size and material.
15. Copy and paste the following code, and define the final time point of interest as the variable “tf_transfection”.

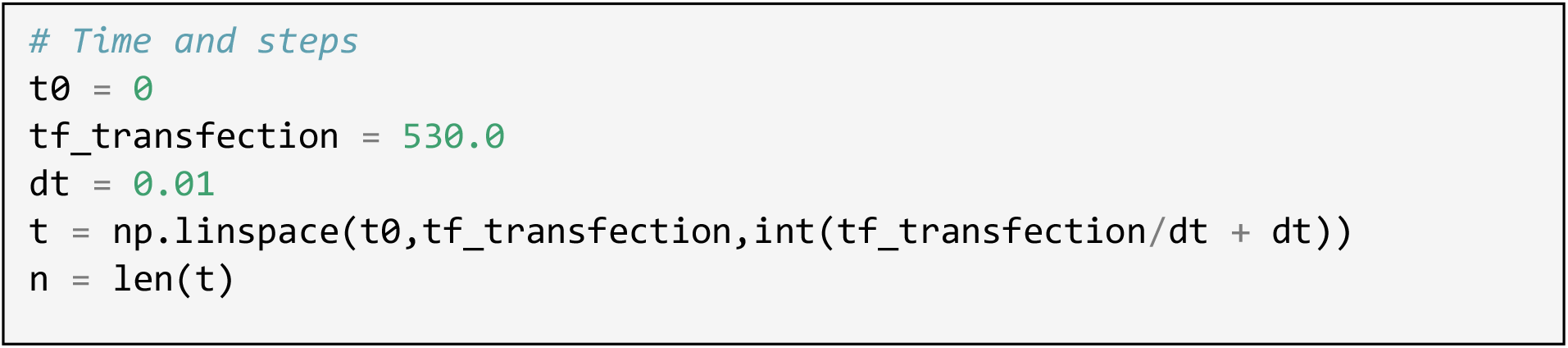
16. Copy and paste the following code, and input the internalization rate fitted using the previous

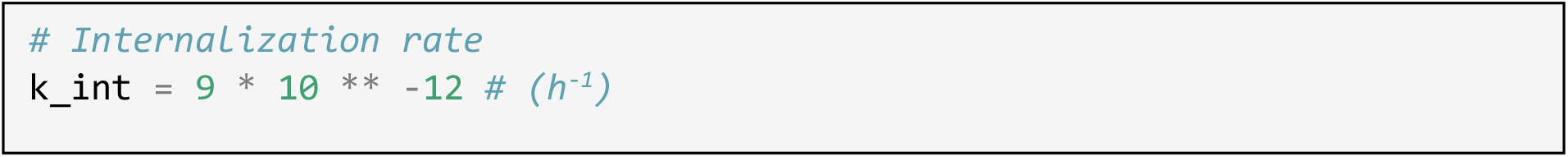

procedure in the ‘Fitting for internalization rate’ section. ***Note:*** Make sure that the internalization rate is the same as the optimized rate obtained in the ‘Fitting for internalization rate’ section. The example code given here includes a different internalization rate value from the previous section because the *in vivo* example is modeling a different experimental scenario.
17. Copy and paste the following code, and vary the clearance rate.

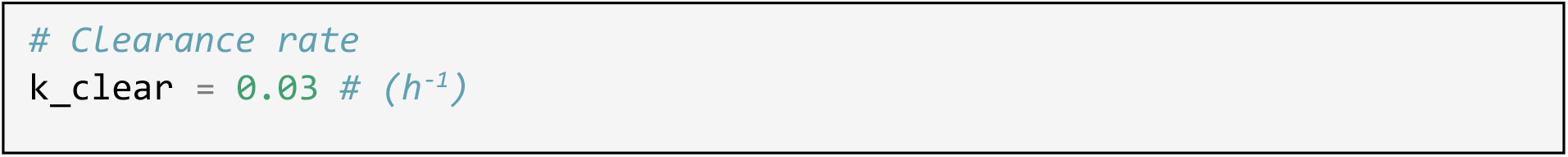 ***Note:*** The clearance rate is used to correctly depict the amount of external siRNA over time. Exact clearance rates for different tissues are not as widely characterized experimentally, and users may have to vary this rate manually until a good alignment is achieved between experimental and modeling data.
18. Copy and paste the following code, run code, and obtain changes in mRNA and protein concentration levels.

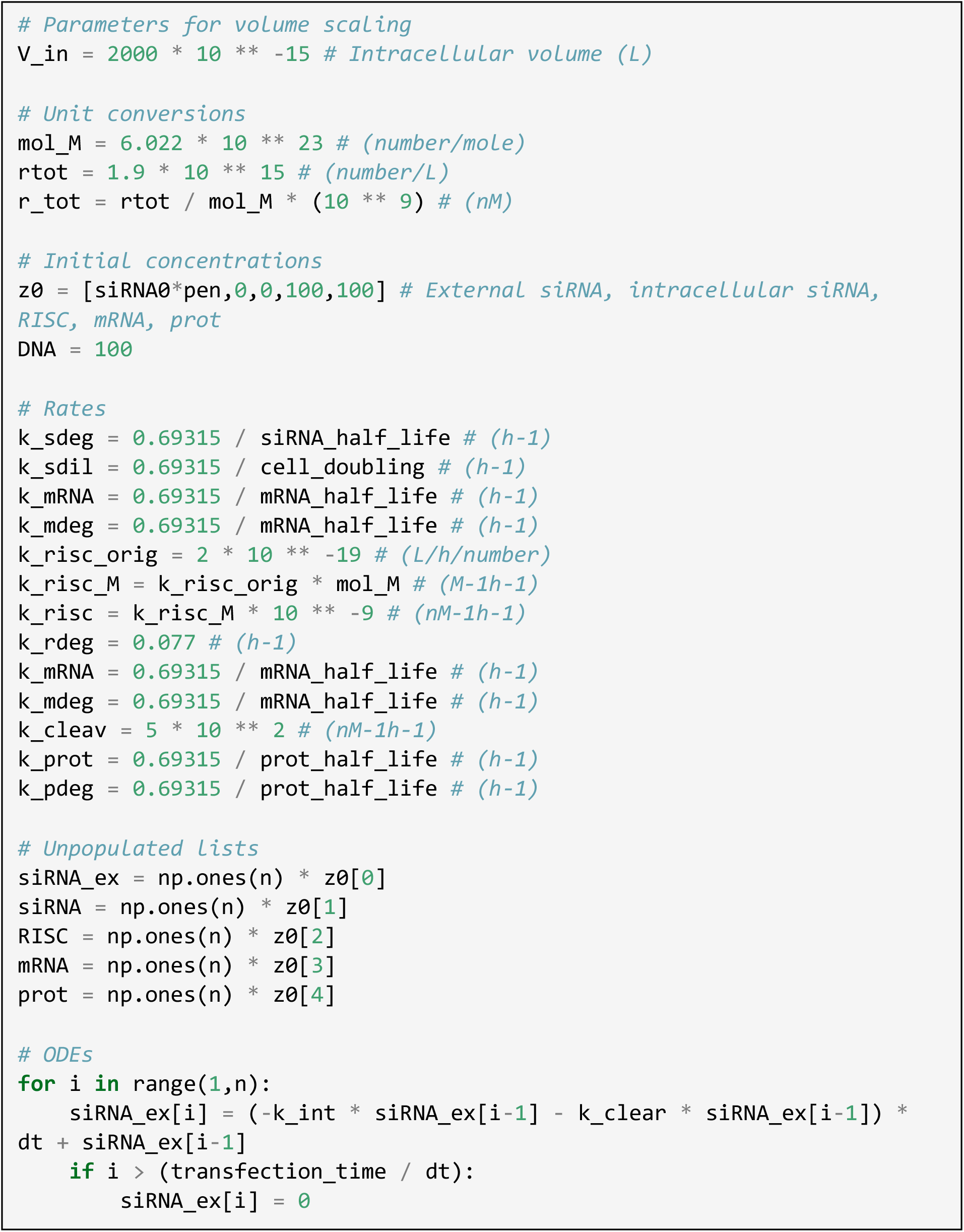
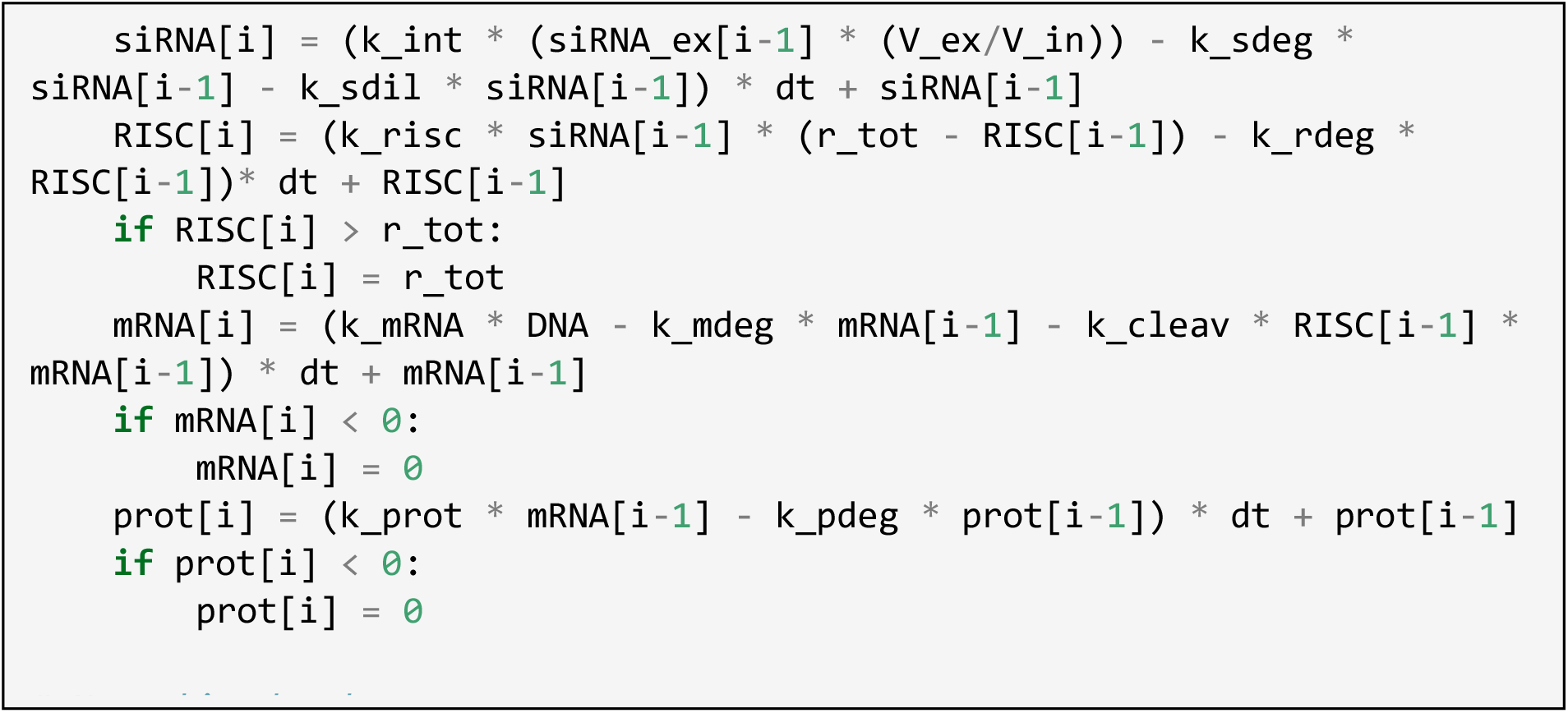
19. Copy and paste the following code and plot desired data accordingly.

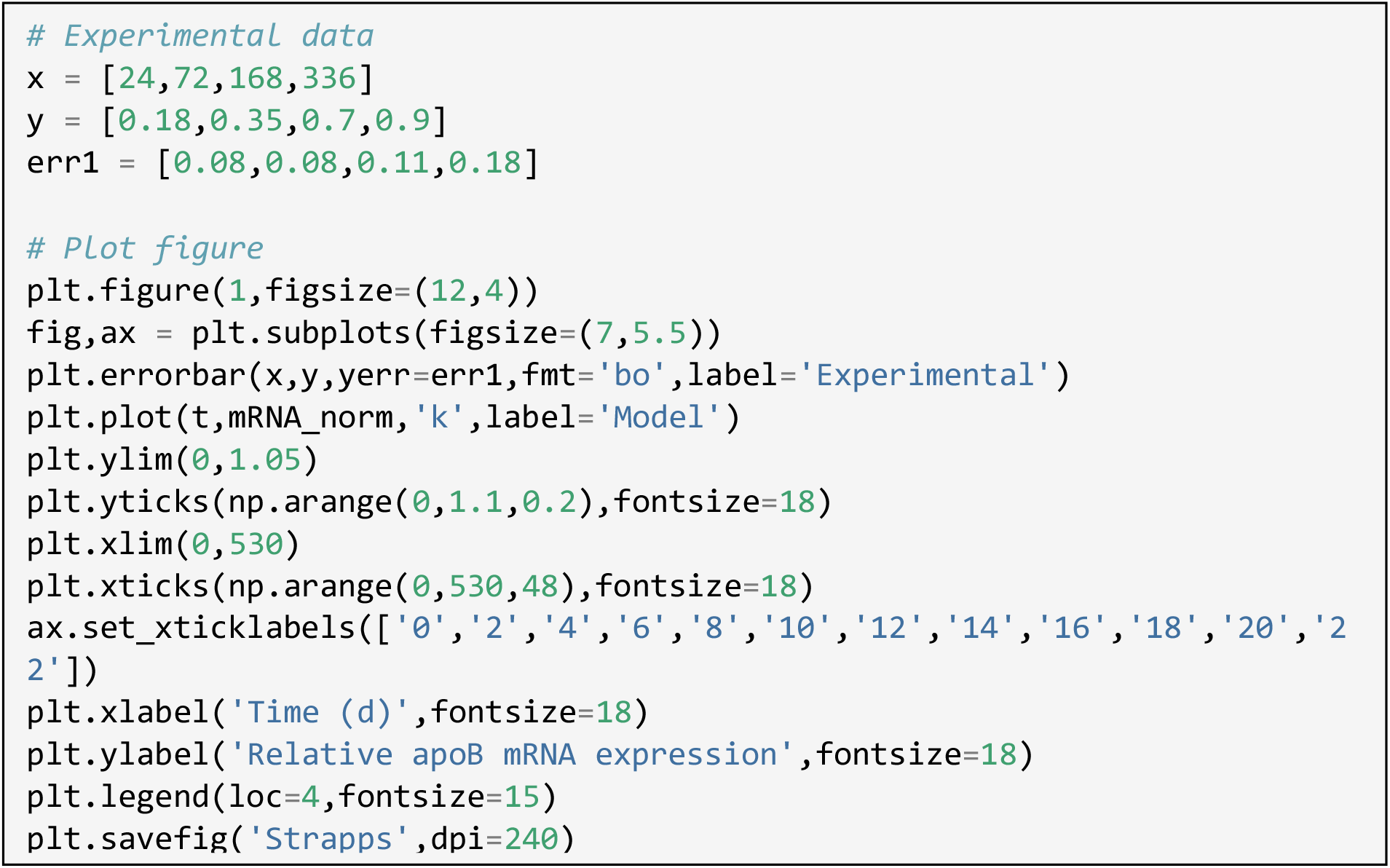 ***Note:*** In this example code, experimental data regarding apolipoprotein B expression levels over time are incorporated within the code. Users can change data within the x-, y-, and err1-arrays for time points, gene expression levels, and associated error, respectively. The ‘# Experimental data’ section of the code can be omitted if users do not wish to plot experimental and modeling data together. Also, plot details including plot type, x-axis and y-axis ticks, and x-axis and y-axis limits can be customized to meet the users’ needs. Detailed explanations and examples can be found at matplotlib.org.(Hunter, 2007)

**Figure 4.**
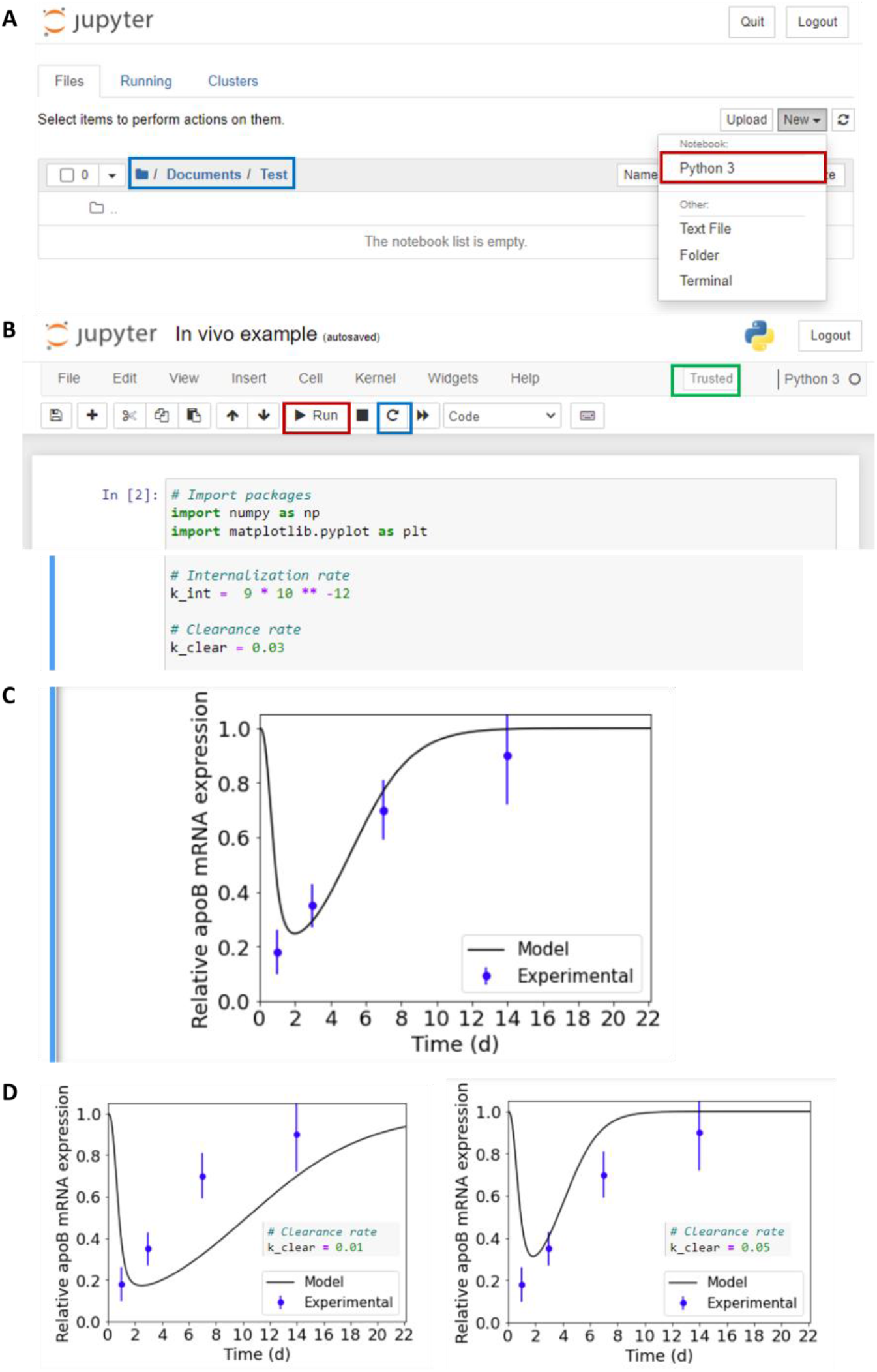
Modeling changes in gene expression levels *in vivo* using the fitted internalization rate. (A) As mentioned in previous section, after opening Jupyter Notebook, choose the correct directory in which to save your notebook (blue box). Click on the ‘New’ box and create a new Python 3 notebook (red box). (B) Copy and paste all code provided in the ‘Modeling siRNA-induced changes in gene expression levels *in vivo*’ section. Make sure that the internalization rate is the same as the optimized rate obtained in the ‘Fitting for internalization rate’ section. If there is an error and users must re-run the code for any reason, make sure to reset the kernel (blue box) before re-running. Make sure that the kernel is trusted (green box). Input an initial guess for the clearance rate. (C) Once the code is finished, a plot will be generated with the details and data provided in the ‘# Plot figure’ section of the code. (D) Vary the clearance rate until model results and experimental data are in good agreement.

## Expected outcomes

### Determining a reasonable internalization rate

By running the code provided in ‘Fitting for internalization rate,’ users will obtain an internalization rate that results in the lowest RMSE when using the experimental data point(s) defined in step 4. In an earlier publication (Roh et al., 2021) and as shown in Figure 5, we have validated that an internalization rate fitted from an *in vitro* experimental measurement of gene expression at a single time point produced predictions for gene silencing efficiencies that were in remarkable agreement with the predictions using an internalization rate fitted from experimental measurements of gene expression at multiple time points.

**Figure 5.**
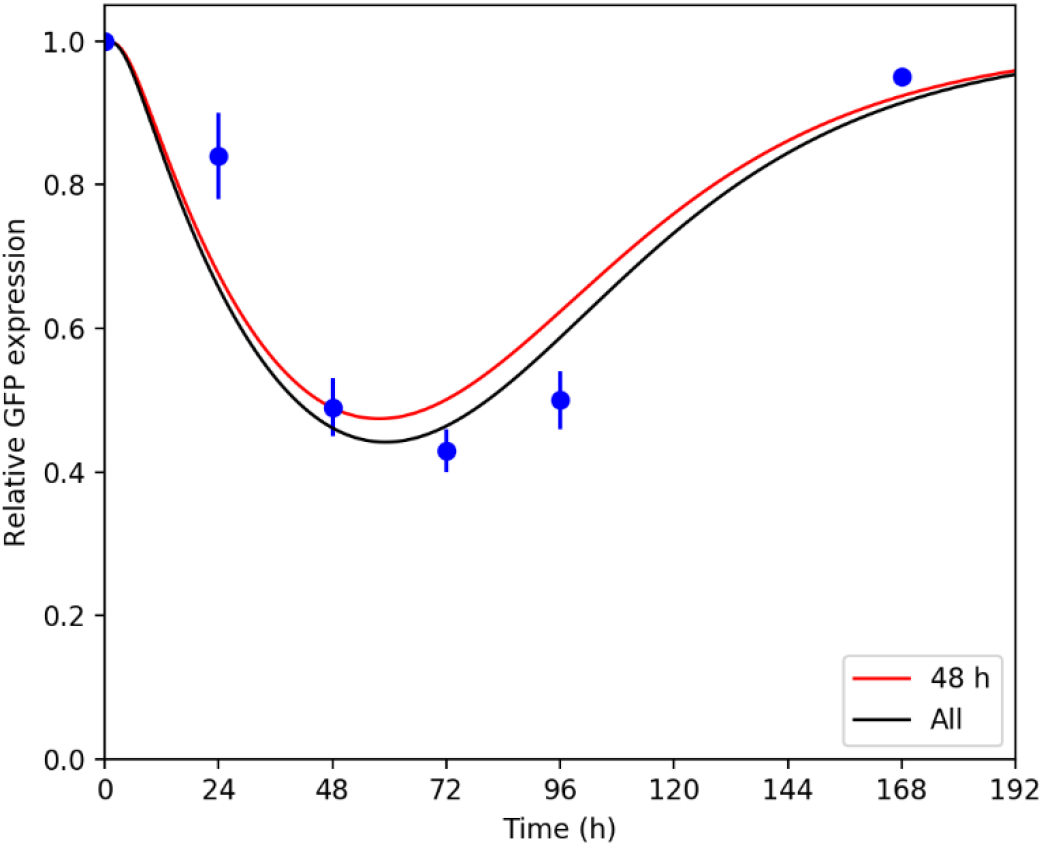
Comparison of model predictions using a single-time-point fit vs. multiple-time-point fit. GFP expression kinetics modulated by naked siRNA *via* electrotransfection in B16F10 cells predicted by the kinetic model using either a single-time-point-fit at 48 h (red line) or a multiple-time-point fit (black line) plotted in conjunction with experimental data (blue dots). All data points and error bars for standard deviations were generated using data reported by the original authors.(Paganin-Gioanni et al., 2011, Roh et al., 2021)

For this procedure, we recommend using a time point between 48 and 72 h, if possible, as mentioned in the Troubleshooting and Limitations sections. This time point is appropriate because (1) it is not short enough to run into the limitations associated with our definition of internalization rate as an aggregated parameter of multiple steps; (2) it is long enough to reflect changes in protein concentrations; and (3) it is not too long so that protein expressions start recovering back to normal expression levels. An example of model predictions obtained from a single-time-point fit of time points ranging from 24 to 168 h is shown in Figure 6.

**Figure 6.**
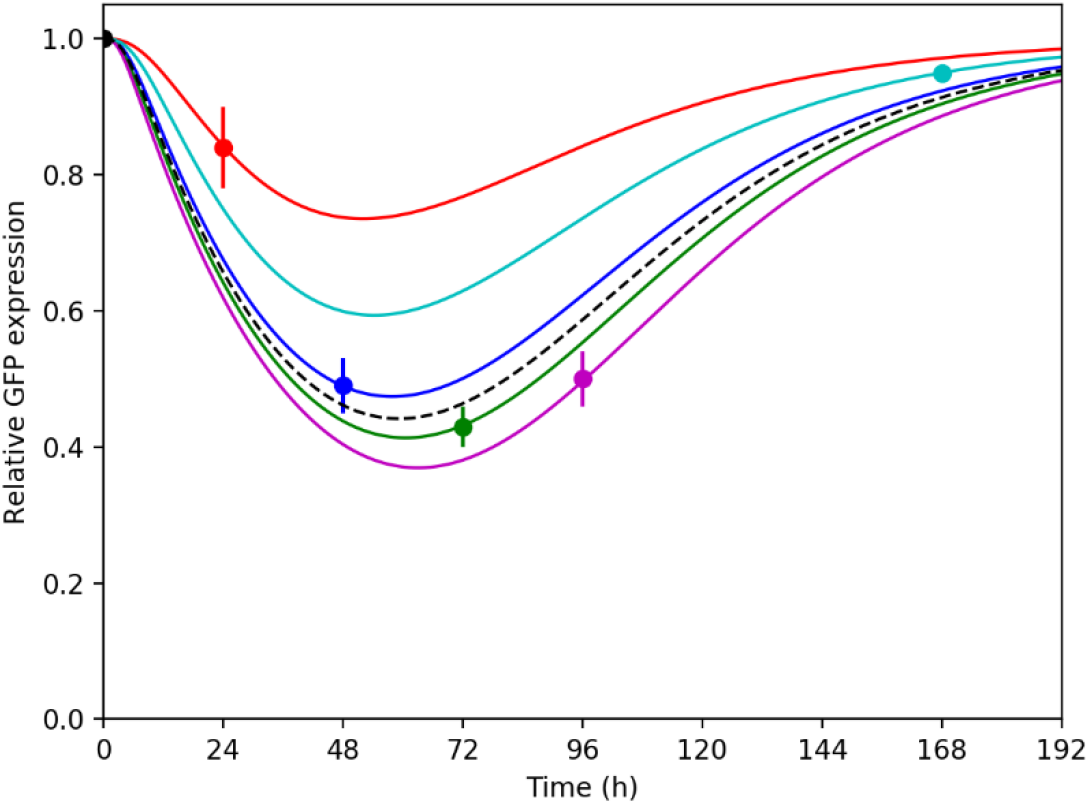
Comparison of model predictions using a single-time-point fit of different time points. GFP expression kinetics modulated by naked siRNA *via* electrotransfection in B16F10 cells predicted by the kinetic model using a single-time-point-fit from experimental data collected at 24 h (red), 48 h (blue), 72 h (green), 96 h (magenta), and 168 h (cyan) compared to results using all five experimental data points to obtain an internalization rate (dotted black line). All data points and error bars for standard deviations were generated using data reported by the original authors.(Paganin-Gioanni et al., 2011, Roh et al., 2021)

### Generating model predictions for siRNA formulation efficacy *in vitro* and *in vivo*

If a reasonable internalization rate was obtained using the above procedure, users will be able to get good agreement between model predictions and experimental data by running the code provided in ‘Modeling siRNA-induced changes in gene expression levels *in vitro*.’ An example result is shown in Figure 7.

**Figure 7.**
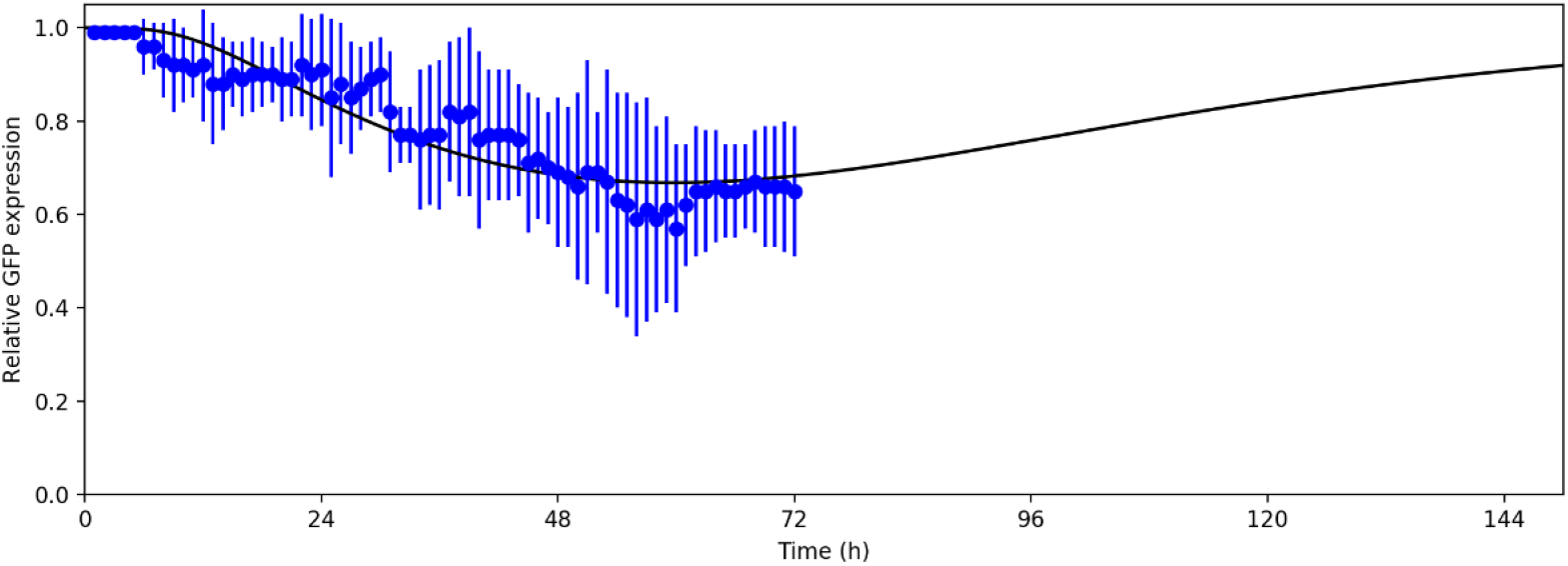
Comparison of model predictions and experimental results of *in vitro* GFP silencing. Model prediction (black) compared to experimental values (blue) for changes in *in vitro* protein expression levels of GFP in HeLa cells using calcium phosphate nanoparticles. All data points and error bars for standard deviations were generated using data reported by the original authors.(Chernousova and Epple, 2017, Roh et al., 2021)

For the *in vivo* modeling, users will be required to vary tissue clearance rates manually because these rates are not widely available in literature. Example results of how model predictions change with different clearance rates are shown in Figure 8.

**Figure 8.**
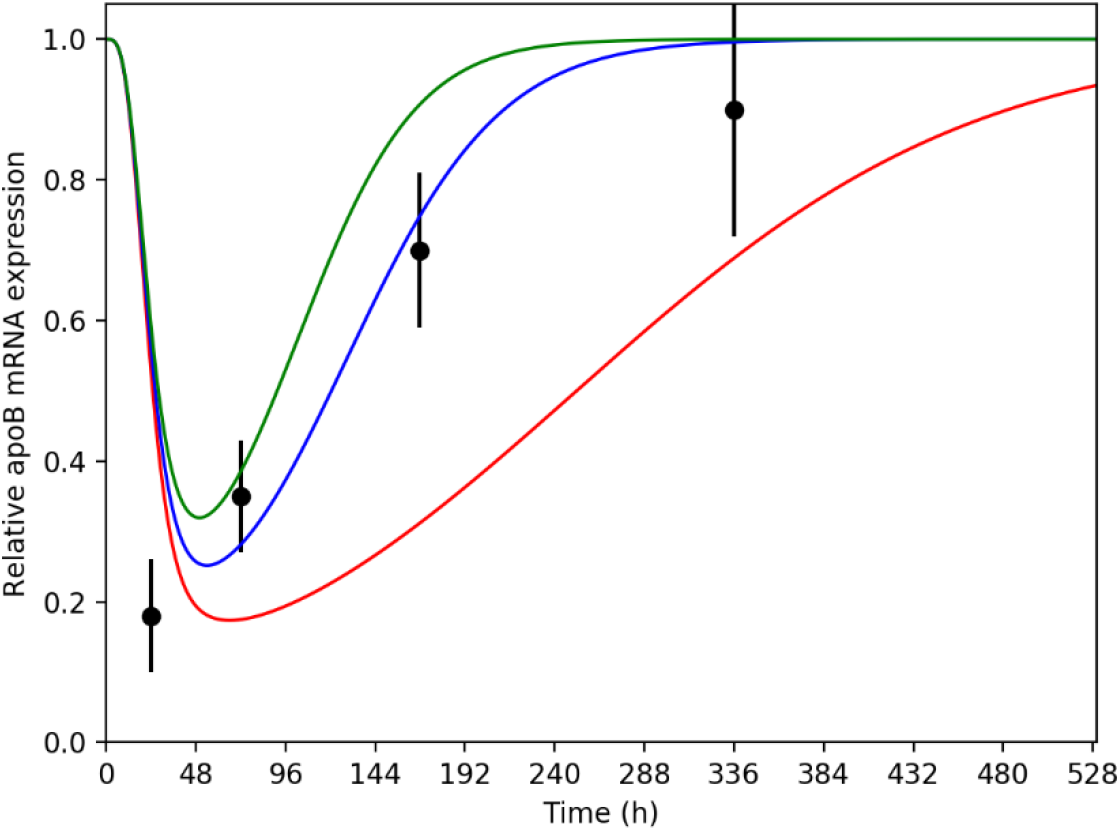
Comparison of model predictions using different clearance rates and experimental results of *in vivo* apoB silencing. *In vivo* model predictions (green line, clearance rate = 0.05 h^-1^; blue line, clearance rate = 0.03 h^-1^; red line, clearance rate = 0.01 h^-1^) compared to *in vivo* experimental values (black) of mRNA expression levels in responses to a single siRNA dose over time when delivered in the form of a siRNA-polymer conjugate. All data points and error bars for standard deviations were generated using data reported by the original authors.(Strapps et al., 2010, Roh et al., 2021)

## Limitations

Due to the assumptions made throughout the model, there is some limitation to the single-time-point fit approach. Specifically, we suspect that there are limitations to obtaining accurate fits for the single unknown parameter, the internalization rate, using earlier time points because we are aggregating multiple steps that are hard to characterize for the internalization process. For reasonable model predictions, users should select a time point that is consistent with the expected kinetics of the comprehensive internalization process. A suggested time point would be 48 h post-siRNA exposure because this time point is longer than most protein half-lives (*i*.*e*., a time point that would reflect changes in protein concentrations after siRNA internalization).

## Troubleshooting

### Problem 1

Optimization takes a very long time to complete (during internalization rate fitting, <u>step 6</u>)

### Potential solution

A possible reason for this problem is the initial guess was far from the optimal value or an excessive number of experimental time points was specified. Update the internalization rate value (recommended starting value is 10^−10^ h^-1^) or reduce the number of experimental measurement inputs.

### Problem 2

Model results do not align with experimental data using optimized internalization rate based on fitting data from a single time point (during *in vitro* modeling, <u>steps 7-11</u>)

### Potential solution

As discussed in the ‘Limitations’ section, using experimental data taken at too early of a time point can result in inaccurate fits for the internalization rate. Choose a time point between 48 and 72 h.

## Resource availability

### Lead contact

Further information and requests for resources and reagents should be directed to and will be fulfilled by the lead contact, Thomas H. Epps, III.

### Materials availability

This study did not generate new unique reagents.

### Data and code availability

The published article includes all datasets/code generated or analyzed during this study.

## Acknowledgments

This work was funded in part by the National Institutes of Health (T32 GM133395 and R21 EB028108-02) and in part by the National Science Foundation (NSF BMAT 1507540). The contents of the manuscript do not necessarily reflect the views of the funding agencies.

## Author contributions

Conceptualization, E.H.R, M.O.S., and T.H.E.; Methodology, E.H.R., M.O.S., and T.H.E.; Investigation, E.H.R.; Resources, M.O.S. and T.H.E.; Writing—Original Draft, E.H.R., M.O.S., and T.H.E; Writing—Review & Editing, E.H.R., M.O.S., and T.H.E.; Visualization, E.H.R.; Supervision, M.O.S. and T.H.E.; Funding Acquisition, M.O.S. and T.H.E.

## References

Alexis, F., Pridgen, E., Molnar, L. K. & Farokhzad, O. C. 2008. Factors affecting the clearance and biodistribution of polymeric nanoparticles. Mol. Pharmaceutics, 5, 505–515.

Bartlett, D. W. & Davis, M. E. 2006. Insights into the kinetics of siRNA-mediated gene silencing from live-cell and live-animal bioluminescent imaging. Nucleic Acids Res, 34, 322–33.

Blanco, E., Shen, H. & Ferrari, M. 2015. Principles of nanoparticle design for overcoming biological barriers to drug delivery. Nat. Biotechnol., 33, 941–951.

Chernousova, S. & Epple, M. 2017. Live-cell imaging to compare the transfection and gene silencing efficiency of calcium phosphate nanoparticles and a liposomal transfection agent. Gene Therapy, 24, 282–289.

Cuccato, G., Polynikis, A., Siciliano, V., Graziano, M., Di Bernardo, M. & Di Bernardo, D. 2011. Modeling RNA interference in mammalian cells. BMC Systems Biology, 5, 19.

Haley, B. & Zamore, P. D. 2004. Kinetic analysis of the RNAi enzyme complex. Nature Structural & Molecular Biology, 11, 599–606.

Harris, C. R., Millman, K. J., Van Der Walt, S. J., Gommers, R., Virtanen, P., Cournapeau, D., Wieser, E., Taylor, J., Berg, S., Smith, N. J., Kern, R., Picus, M., Hoyer, S., Van Kerkwijk, M. H., Brett, M., Haldane, A., Del Río, J. F., Wiebe, M., Peterson, P., GÉrard-Marchant, P., Sheppard, K., Reddy, T., Weckesser, W., Abbasi, H., Gohlke, C. & Oliphant, T. E. 2020. Array programming with NumPy. Nature, 585, 357–362.

He, C., Hu, Y., Yin, L., Tang, C. & Yin, C. 2010. Effects of particle size and surface charge on cellular uptake and biodistribution of polymeric nanoparticles. Biomaterials, 31, 3657–3666.

Hoshyar, N., Gray, S., Han, H. & Bao, G. 2016. The effect of nanoparticle size on in vivo pharmacokinetics and cellular interaction. Nanomedicine, 11, 673–692.

Hunter, J. D. 2007. Matplotlib: A 2D graphics environment. Computing in Science & Engineering, 9, 90–95.

Jasinski, D. L., Li, H. & Guo, P. 2018. The effect of size and shape of RNA nanoparticles on biodistribution. Mol. Ther., 26, 784–792.

Oraiopoulou, M. E., Tzamali, E., Tzedakis, G., Vakis, A., Papamatheakis, J. & Sakkalis, V. 2017. In vitro/in silico study on the role of doubling time heterogeneity among primary glioblastoma cell lines. BioMed Res. Int., 2017, 8569328.

Paganin-Gioanni, A., Bellard, E., Escoffre, J. M., Rols, M. P., TeissiÉ, J. & Golzio, M. 2011. Direct visualization at the single-cell level of siRNA electrotransfer into cancer cells. Proceedings of the National Academy of Sciences, 108, 10443–10447.

Pedregosa, F., Varoquaux, G., Gramfort, A., Michel, V., Thirion, B., Grisel, O., Blondel, M., Prettenhofer, P., Weiss, R., Dubourg, V., Vanderplas, J., Passos, A., Cournapeau, D., Brucher, M., Perrot, M. & Duchesnay, É. 2011. Scikit-learn: Machine Learning in Python. J. Mach. Learn. Res., 12, 2825–2830.

PÉrez-Campaña, C., Gómez-Vallejo, V., Puigivila, M., Martín, A., Calvo-Fernández, T., Moya, S. E., Ziolo, R. F., Reese, T. & Llop, J. 2013. Biodistribution of different sized nanoparticles assessed by positron emission tomography: A general strategy for direct activation of metal oxide particles. ACS Nano, 7, 3498–3505.

Pratt, A. J. & Macrae, I. J. 2009. The RNA-induced silencing complex: a versatile gene-silencing machine. The Journal of biological chemistry, 284, 17897–17901.

Roh, E. H., Epps, T. H. & Sullivan, M. O. 2021. Kinetic modeling to accelerate the development of nucleic acid formulations. ACS Nano, 15, 16055–16066.

Sakurai, K., Chomchan, P. & Rossi, J. J. 2010. Silencing of gene expression in cultured cells using small interfering RNAs. Curr Protoc Cell Biol, Chapter 27, Unit 27.1.1-28.

Strapps, W. R., Pickering, V., Muiru, G. T., Rice, J., Orsborn, S., Polisky, B. A., Sachs, A. & Bartz, S. R. 2010. The siRNA sequence and guide strand overhangs are determinants of in vivo duration of silencing. Nucleic Acids Research, 38, 4788–4797.

Virtanen, P., Gommers, R., Oliphant, T. E., Haberland, M., Reddy, T., Cournapeau, D., Burovski, E., Peterson, P., Weckesser, W., Bright, J., Van Der Walt, S. J., Brett, M., Wilson, J., Millman, K. J., Mayorov, N., Nelson, A. R. J., Jones, E., Kern, R., Larson, E., Carey, C. J., Polat, İ., Feng, Y., Moore, E. W., Vanderplas, J., Laxalde, D., Perktold, J., Cimrman, R., Henriksen, I., Quintero, E. A., Harris, C. R., Archibald, A. M., Ribeiro, A. H., Pedregosa, F., Van Mulbregt, P., Vijaykumar, A., Bardelli, A. P., Rothberg, A., Hilboll, A., Kloeckner, A., Scopatz, A., Lee, A., Rokem, A., Woods, C. N., Fulton, C., Masson, C., HÄggstrÖm, C., Fitzgerald, C., Nicholson, D. A., Hagen, D. R., Pasechnik, D. V., Olivetti, E., Martin, E., Wieser, E., Silva, F., Lenders, F., Wilhelm, F., Young, G., Price, G. A., Ingold, G.-L., Allen, G. E., Lee, G. R., Audren, H., Probst, I., Dietrich, J. P., Silterra, J., Webber, J. T., Slavič, J., Nothman, J., Buchner, J., Kulick, J., SchÖnberger, J. L., De Miranda Cardoso, J. V., Reimer, J., Harrington, J., Rodríguez, J. L. C., Nunez-Iglesias, J., Kuczynski, J., Tritz, K., Thoma, M., Newville, M., Kümmerer, M., Bolingbroke, M., Tartre, M., Pak, M., Smith, N. J., Nowaczyk, N., Shebanov, N., Pavlyk, O., Brodtkorb, P. A., Lee, P., Mcgibbon, R. T., Feldbauer, R., Lewis, S., Tygier, S., Sievert, S., Vigna, S., Peterson, S., More, S., Pudlik, T., Oshima, T., et al. 2020. SciPy 1.0: fundamental algorithms for scientific computing in Python. Nature Methods, 17, 261–272.

Wittrup, A., Ai, A., Liu, X., Hamar, P., Trifonova, R., Charisse, K., Manoharan, M., Kirchhausen, T. & Lieberman, J. 2015. Visualizing lipid-formulated siRNA release from endosomes and target gene knockdown. Nat Biotechnol, 33, 870–6.

